# X chromosome inactivation in the human placenta is patchy and distinct from adult tissues

**DOI:** 10.1101/785105

**Authors:** Tanya N. Phung, Kimberly C. Olney, Michelle Silasi, Lauren Perley, Jane O’Bryan, Harvey J. Kliman, Melissa A. Wilson

## Abstract

One of the X chromosomes in genetic females is silenced by a process called X chromosome inactivation (XCI). Variation in XCI across the placenta may contribute to observed sex differences and variability in pregnancy outcomes. However, XCI has predominantly been studied in human adult tissues. Here we sequenced and analyzed DNA and RNA from two locations from 30 full-term pregnancies. Implementing an allele specific approach to examine XCI, we report evidence that XCI in the human placenta is patchy, with large patches of either silenced maternal or paternal X chromosomes. Further, using similar measurements, we show that this is in contrast to adult tissues, which generally exhibit mosaic X-inactivation, where bulk samples exhibit both maternal and paternal X chromosome expression. Further, by comparing skewed samples in placenta and adult tissues, we identify genes that are uniquely silenced or expressed in the placenta compared to adult tissues highlighting the need for tissue-specific maps of XCI.

## Introduction

X chromosome inactivation (XCI) evolved in eutherian mammals, at the same time as the development of invasive placentation, and is important for many biological processes (Natri et al. 2019). XCI is a mechanism to regulate the dosage of gene expression due to the differences in the number of X chromosomes between the sexes: genetic females (XX) have two X chromosomes while genetic males (XY) have only one X chromosome (Lyon 1961; Wilson Sayres and Makova 2013).

There are two levels of XCI. The first level is at the whole chromosome level, where either the maternal X or the paternal X is chosen for inactivation. The choice of which X is inactivated has been observed to happen in one of two ways: random XCI (the maternal and paternal X are silenced with equal probability), and imprinted XCI (the paternal X is inactivated). In mice, the extraembryonic lineages that ultimately give rise to the placenta and some extraembryonic membranes show paternally imprinted XCI (Takagi and Sasaki 1975; K. D. Huynh and Lee 2001; Khanh D. Huynh and Lee 2005). Paternally imprinted XCI has also been reported in rats (Wake, Takagi, and Sasaki 1976), cows (Xue et al. 2002), and marsupial mammals (Richardson, Czuppon, and Sharman 1971; Cooper et al. 1989; Al Nadaf et al. 2010). However, in mice, the embryonic lineages that ultimately give rise to the rest of the fetus exhibit random XCI (Takagi and Sasaki 1975). Random XCI has also been reported in mule and horse placenta (Wang et al. 2012).

The second level of XCI concerns which individual genes are subject to inactivation (silenced or not expressed) on the inactive X chromosome. Previous studies have aimed to determine what genes on the X chromosome escape XCI in a variety of tissues (see (Carrel and Brown 2017) for an overview of expression from the inactive X). Carrel and Willard (2005) studied biallelic expression in primary human fibroblast cell lines and rodent/human somatic hybrids (Carrel and Willard 2005). Cotton et al. (2015) used DNA methylation data to characterize escape status in 27 adult tissues (Cotton et al. 2015). Gene-specific silencing has also been shown to vary between tissues and individuals in mouse and human (Berletch et al. 2015; Tukiainen et al. 2017). However, previous cross-tissue studies on gene-specific escape from XCI did not include the placenta.

The human placenta is a transient tissue that is formed early in pregnancy and develops from the outer layer of the pre-implantation embryos (James, Carter, and Chamley 2012; Turco and Moffett 2019). The placenta has the genotype of the fetus and plays a critical role in pregnancy by regulating nutrition and protecting the developing fetus from the pregnant person’s immune system (Gude et al. 2004). Improper placenta development can lead to complications such as preeclampsia and fetal growth restriction (Rathbun and Hildebrand 2019). It is important to study gene-specific escape in the placenta because XCI could play an important role in pregnancy complications; Gong et al. (2018) found that the spermine synthase gene (SMS) that modulates fetal growth restriction, a common pregnancy complication may escape XCI in the placenta but not in other tissues (Gong et al. 2018). Further, the degree of XCI skewing could be associated with pregnancy loss (Sui, Chen, and Sun 2015). Therefore, understanding XCI in this early-formed tissue is critical for understanding downstream developmental effects.

Previous research suggests that XCI across the entire X chromosome in the placenta is random and patchy, but these studies relied on a limited number of loci, and did not investigate chromosome-wide gene-specific variability (Moreira de Mello et al. 2010; Peñaherrera et al. 2012). Further, one of the challenges of previous research on XCI is that indirect methods have been used to infer inactivation. Gong et al. (2018) and Tukiainene et al. (2017) used higher expression in females as compared to males as a proxy for escape genes, rather than directly measuring inactivation status. However, Slavney et al. (2016) observed that escape genes show higher expression compared to inactivated genes in both males and females (Slavney et al. 2016).

Here we utilized next-generation sequencing to characterize genome-wide patterns of XCI in the human placenta, to compare patterns of XCI between the placenta and other tissues in humans, and to analyze genes that escape XCI that are unique to the placenta. We examined allele specific expression by sequencing whole-exome and whole-transcriptome (from two locations) of 30 full-term placentas from uncomplicated pregnancies. We further utilized data from the Genotype-Tissue Expression (GTEx) projects to examine XCI patterns in adult tissues (GTEx Consortium 2020). Using our experimental approach, we have the capacity to observe: 1) extreme skewing in the human placenta, where all placentas show the same X inactivated in both sites; 2) patchiness, where the placenta exhibits patches of daughter cells either maternal or paternal X expression only; or, 3) mosaicism, where clusters of cells in a sample exhibit both maternal and paternal X chromosome expression (**Figure 1**). Here we limit analysis of gene-specific inactivation to placental and adult tissues with skewed XCI to directly measure allele-specific expression. We observed evidence for skewing (large patches of exclusively maternal or paternal X expression) in the human placenta that is not present in adult tissues. We also compared genes that are inactivated, genes that escape from inactivation, and genes that show variable escape between the placenta and adult tissues. While a majority of genes are concordant for inactivation status between the placenta and adult tissues (71%), we observed a subset of genes that show opposite patterns between the placenta and adult tissues.

**Figure 1.**
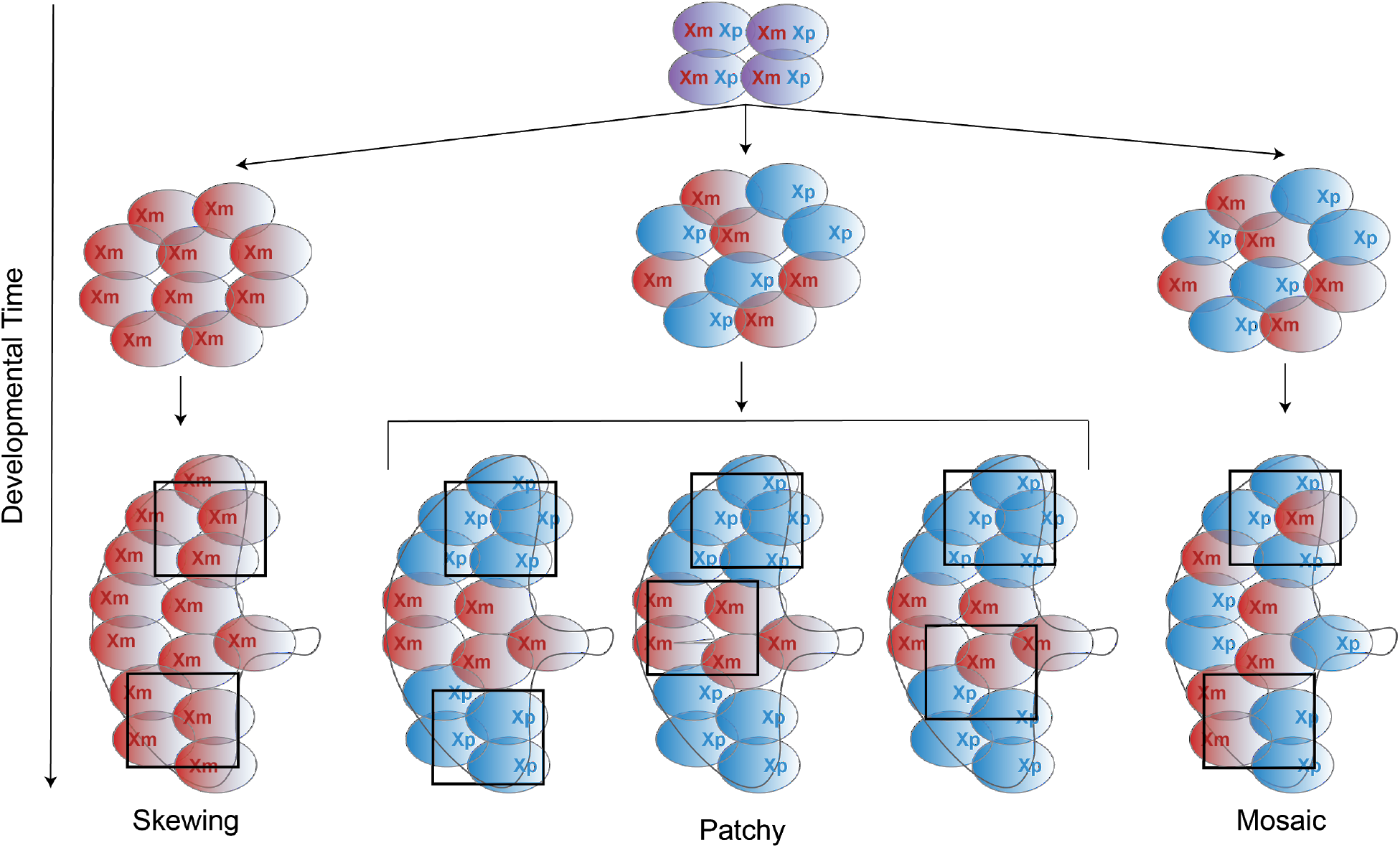
Schematic possible patterns of X chromosome inactivation. Prior to XCI, both the maternal and the paternal X chromosomes are present (purple cells). The blue cell represents the maternal X is inactivated, and the red cell represents the paternal X is inactivated. If XCI occurs very early in development and propagates to all daughter cells, we would observe extreme skewing where all placentas show the same X inactivated. In this case, the same X chromosome is inactivated in both sites sampled from this study. If XCI occurs at an intermediate stage, XCI is random but is present with large patches of daughter cells with only the maternal X or paternal X. In this case, three possible scenarios can happen: 1. Both of the sites sampled in this study happen to come from two patches with the same X inactivated, 2. Each site sampled in this study comes from two patches with different X inactivation, and 3. One of the sites sampled in this study comes from the boundary of two patches. If XCI occurs very late in development, there are either small or no patches, creating a mosaic pattern of XCI.

Our results provide additional evidence that XCI in the human placenta is random and patchy. Our study further shows that patterns of X-inactivation differ between embryonic tissue and adult tissues, consistent with different developmental structures of these tissues. Further, we identified genes with unique XCI status in the placenta. We hypothesize that these genes could play an important role in the placenta and pregnancy.

## Results

### Whole chromosome level of X chromosome inactivation differs between the placenta and adult tissues

We observed evidence for patchiness in XCI in the placenta. To examine patterns of XCI in the human placenta, we sequenced the whole transcriptome from two separate extraction sites of the same placenta. We then employed a phasing strategy on the X chromosome using whole exome and whole transcriptome sequence data. Briefly, at heterozygous sites that exhibit allelic imbalance, we reasoned that the biased alleles (the alleles that are expressed in a higher proportion) are all on the same X chromosome (see **Methods**). In 17/29 placentas, both sites in the placenta exhibited extremely skewed X-inactivation, either with both extraction sites showing the same inactivated X chromosome (12 placentas; **Figure 2A**; **Figure S7A**) or each site showing the opposite X chromosome inactivated (5 placentas; **Figure 2A**; **Figure S7B**). In the remaining twelve placentas we observed one extraction site showing skewed XCI, and the other showing variable proportions of both X chromosomes being expressed (**Figure 2A**; **Figure S7C**). In this subset, we postulated that one of our samples was collected on the boundary of two different patches of X inactivated cells. To validate that the patterns we observed on the X chromosome are indeed XCI, we repeated the same analyses on chromosome 8, where all samples showed biallelic expression (**Figure 2B**). Our results suggest that the human placenta is organized into large patches with either the maternal or paternal X chromosome being inactivated.

**Figure 2.**
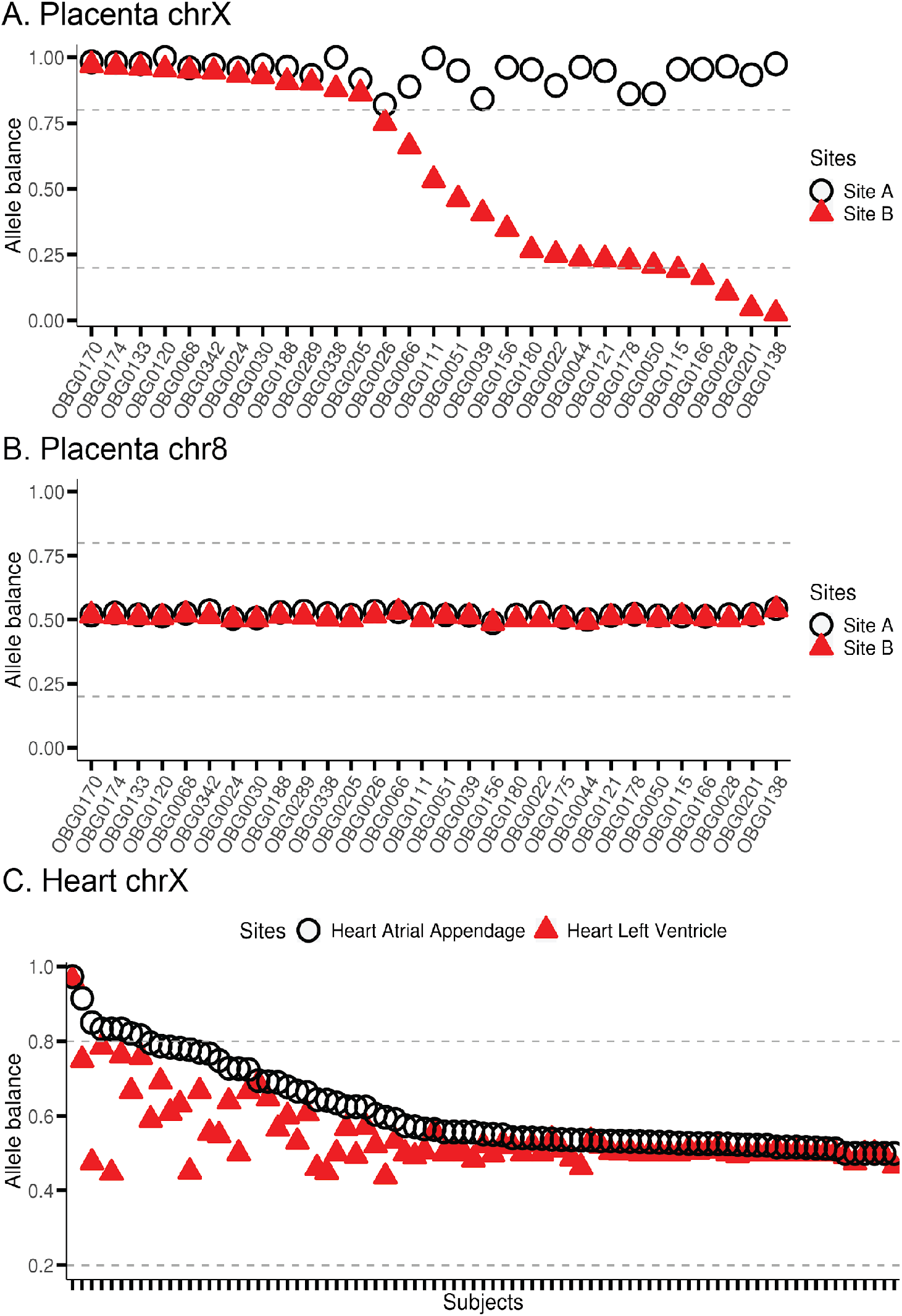
X chromosome inactivation is patchy in the human placenta and mosaic in the human heart. Phased allele balance aggregated across variants for each sample is shown for site A (open black circle) and site B (filled red triangle). Each point is the median value across variants. Median was used to minimize the effect of outliers. The dotted gray horizontal lines denote allele balance of 0.2 and 0.8. (A) Placenta X chromosome. We removed variants that fall within the pseudoautosomal regions (PARs) prior to computing medians. (B) Placenta chromosome 8. (C) Heart X chromosome.

We then found that patterns fXCI in the placenta are different from adult tissues (**Figure 2, Figure 3**). We first compared the pattern of whole chromosome XCI in the two placenta samples we collected with whole chromosome XCI in two regions collected from adult human hearts (left ventricle and atrial appendages) sampled by the GTEx consortium (GTEx Consortium 2020) (**Figure 2A, 2C**). We identified 85 individuals with RNAseq data for both of these locations in the heart and employed the same phasing approach as in the placenta. In the heart tissue, both regions exhibit biallelic expression (both maternal and paternal X expression) in 91% of individuals (**Figure 2C**). This is in stark contrast to the placenta, where none of individuals exhibit biallelic expression at both regions (**Figure 2A**) .

**Figure 3.**
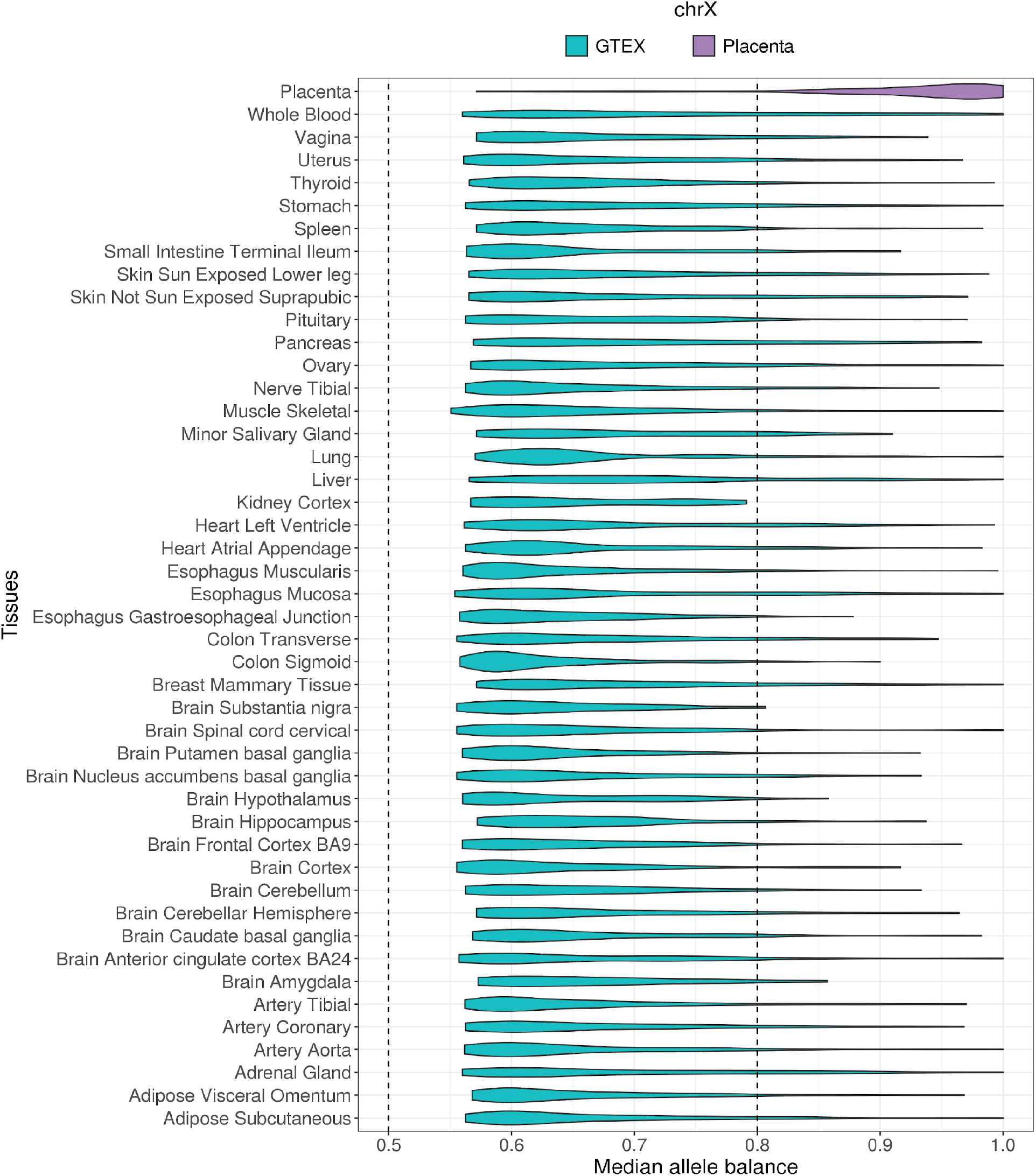
Whole chromosome level of X chromosome inactivation differs between the placenta and adult tissues in humans. Unphased median allele balance is plotted for the placenta in this study (purple) and for 45 adult tissues in the GTEx dataset (blue). Each point of the violin plot is the median allele balance for each sample. Because there are no multiple site samplings for the GTEx data, unphased allele balance was computed. Here, the pseudoautosomal regions of the X chromosome are removed.

We further found the same biallelic pattern of XCI across all 43 adult tissue samples, in contrast to the placenta (**Figure 3**). Because other adult tissues include a single sample (GTEx Consortium 2020), to assess chromosome wide XCI we compared median allele balance computed from the biased allele for each sample. While most placenta samples (90%) exhibit skewed allele balance (*i*.*e*., allele balance greater than 0.8), most samples from adult tissues (89%) exhibit biallelic expression (*i*.*e*., allele balance between 0.5 and 0.8) (**Figure 3**). Similar to the placenta, however, also we observed biallelic expression across all adult tissues on chromosome 8, as expected (**Figure S8**). Together, these results suggest that patterns of whole chromosome XCI differ between the placenta and adult tissues.

### Gene-specific escape comparison between the placenta and adult tissues

We observed a large degree of variability in gene-specific escape in the placenta, both across and within individuals. We categorized genes on the X chromosome using only samples that exhibit skewed median allele balance across the X chromosome (i.e. median allele balance greater than 0.8, see **Methods**). Only 7 genes (2%) show evidence for escape across all samples with the ability to assay, while 73 genes (18%) show evidence of silencing across all samples (**Figure S9**). Rather, most genes, 316 genes (80%), show a gradient of the proportion of samples that exhibit evidence for escape. In addition, heterogeneity exists between two sites within individuals (**Figure S10)**. Given the tremendous heterogeneity in evidence for escape, we additionally investigated the ratio of female-to-male gene expression across X-linked genes. We observed that genes in all three classes (escape, variable, and silenced) exhibit a range of female-to-male gene expression ratios, with higher gene expression in females as compared to males underpowered to determine whether a gene has escaped X-inactivation (**Figure S11**).

We found similarities and differences in gene-specific escape and silencing in the human placenta when compared to the adult tissues. We used 525 skewed adult tissue samples in the GTEx dataset (out of 4958 total samples) and 52 skewed samples in the placenta dataset (out of 58 total samples) to categorize XCI inactivation status (see **Methods**). A gene is considered for this analysis if there are at least five informative samples (i.e., an informative sample is defined as having at least one heterozygous and expressed variant for that gene) in both the placenta dataset and the GTEx dataset. We found 186 such genes. Out of these 186 genes, 132 genes (71%) show the same pattern of escape or silencing between the adult tissues and the placenta dataset (**Figure 4A**). Specifically, 111 genes are inactivated, 15 genes escape, and 6 genes show variable escape between the adult tissues and the placenta. We also observed differences between the adult tissues and the placenta. We found three genes that escape in the placenta but are inactivated in adult tissues: *ACOT9, CASK, TSC22D3* (**Figure 4B**). We found two genes that are inactivated in the placenta but escape in adult tissues: *ARSD* and *PRKX* (**Figure 4C**). We examined the mapping quality and total RNA read counts of the variants in these genes to confirm that these observations are not artifacts (**Figure S12**). These results suggest that while the majority of genes on the X chromosome show consistent XCI between the placenta and adult tissues, a subset exhibits opposite patterns that could be attributed to the unique nature of the placenta tissue, which is an embryonic tissue.

**Figure 4.**
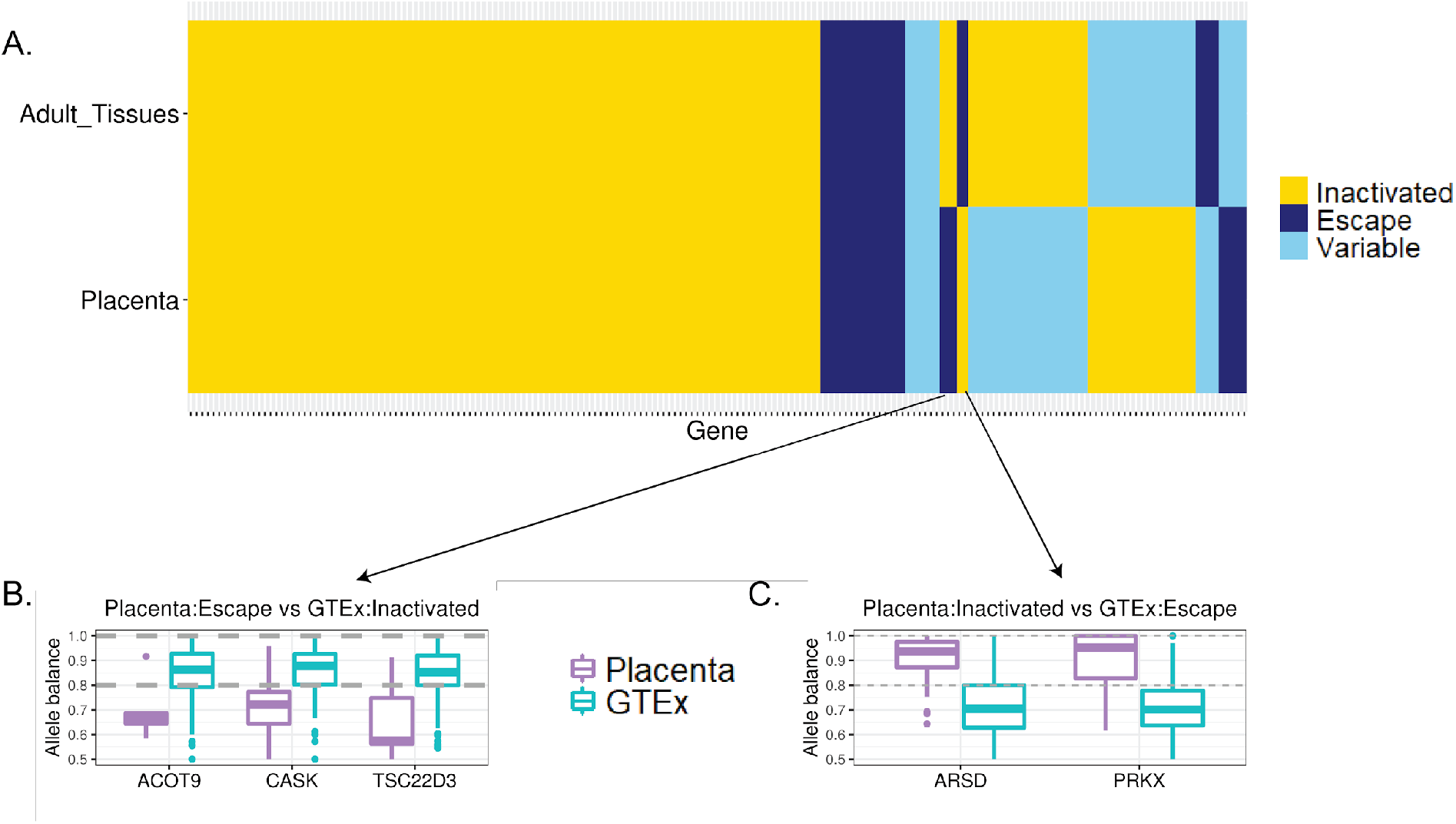
Gene-specific escape comparison between the placenta and adult tissues. (A) A heatmap showing genes that are inactivated in both the placenta and adult GTEx tissues (yellow), genes that escape X chromosome inactivation in both the placenta and adult GTEx tissues (dark blue), and genes that exhibit variable escape across individuals in both the placenta and adult GTEx tissues (light blue). Boxplots showing median allele balance across all samples in the placenta (purple) and adult tissues in GTEx (blue) for the three genes that show evidence for escaping XCI in the placenta but being inactivated in the adult GTEx tissues (B) and for the two genes that show evidence for being inactivated in the placenta but escaping in the adult GTEx tissues (C). Each point of the box plot is the median allele balance computed across variants for that gene (see **Methods**).

## Discussion

We observed that X chromosome inactivation in the human placenta is organized into large patches of maternal or paternal X expression. We utilized whole exome and whole transcriptome sequence data to analyze allele specific expression across female placentas and found evidence that the human placenta exhibits large patches of maternal or paternal X chromosome expression. While patchy XCI in the human placenta has been observed in humans previously, these prior studies rely on a few SNPs and a few genes in a limited number of samples (Moreira de Mello et al. 2010; Looijenga et al. 1999). For example, Looijenga et al. (1999) examined two samples from each of the nine female (46, XX) full term placenta samples, using methylation in a single gene, androgen receptor, as a read-out, they found that three placentas exhibited predominantly maternal X-inactivation, one exhibited paternal X-inactivation, and the remaining 5 exhibited both maternal and paternal alleles (Looijenga et al. 1999). In a different study, with at least 5 informative SNPs used to infer X-inactivation status in a single site collected from 22 placentas, the authors similarly observe some placentas have maternal X-inactivation, some have paternal X-inactivation, and others show biallelic expression across the X-linked SNPs (Moreira de Mello et al. 2010). From this, the authors proposed that the human placenta has large patches of maternal or paternal X-inactivation. These studies typically relied on a single gene as in Looijenga et al. (1999) or did not include multiple samplings from the same placenta as in Moreira de Mello et al. (2010). With improved sampling (i.e. sampling two independent locations from each placenta, from opposing quadrants) and genome-wide sequencing, we confirm that patterns of XCI are predominantly patchy in the human placenta.

We found distinct patterns of XCI between the placenta and adult tissues in humans: while most placenta samples (∼90%) exhibit skewed expression, only about 11% of samples from human adult tissues exhibit skewed expression. This is consistent with previous methylation-based analyses that observed skewed inactivation across different developmental layers of the placenta (Peñaherrera et al. 2012), suggesting that each villus tree is clonally derived from a few precursor cells, and that there is little cell migration across placental regions. This is in contrast to adult tissues, where we observe very little skewing, suggesting more cell migration during development. Though both observations are contingent on the size of the tissue sample analyzed in bulk, where samples with more cells are more likely to capture both X-chromosomes being expressed.

Although we found high concordance in genes that are inactivated and genes that escape between the placenta samples and adult tissues (**Figure 4**), we found a subset of genes with unique XCI status in the placenta. Specifically, we found five genes that show patterns that are unique to the placenta: *ACOT9, CASK*, and *TSC22D3* are inactivated in adult tissues, but escape XCI in the placenta (**Figure 4B**). *ARSD* and *PRKX*, both genes in the pseudoautosomal region, escape XCI in adult tissues but are inactivated consistently across regions in the placenta (**Figure 4C**). *ACOT9* has been associated with syndromic X-linked intellectual disability Turner Type (Stelzer et al. 2016) but the function of *ACOT9* in the placenta is unclear. *CASK* was shown to be subject to XCI in mice and has been implicated to play an important role in neurological disorders (Mori et al. 2019). Since the placenta and cord blood could also be linked to neurological disorders (Mordaunt et al. 2020), it is plausible that *CASK* escapes XCI, or experiences XCI erosion, in the placenta while it is inactivated in other tissues. *TSC22D3* is an immunity related gene. Syrett al. (2019) observed that *TSC22D3* is overexpressed in females compared to males in T cells of patients with systemic lupus erythematosus (Syrett et al. 2019). Curiously, ARSD is active in adult tissues, but silenced in the placenta. Larson et al. (2017) observed that *ARSD* is inactivated in at least one ovarian tumor sample, highlighting a potential parallel between the placenta and tumor development (Larson et al. 2017). *PRKX* is also shown to be an escape gene in adult tissues here, and as previously reported (Wainer Katsir and Linial 2019), but not in the placenta. We hypothesize that these genes could be further studied for a potential role in pregnancy and pregnancy complications.

We additionally show that sex differences in gene expression are not sufficient on their own to determine whether a gene escapes inactivation or not (**Figure S12**). This highlights the importance of understanding both inactivation status, and expression differences between the sexes. Gong et al. (2018) used female-biased gene expression as a proxy for identifying potential genes that escape XCI in the placenta. 28/47 potential escape genes identified by Gong et al. (2018) were reported previously as being inactivated or unknown (Gong et al. 2018). Of these, 3/28 genes exhibit variable escape in our placenta data (*MBTPS2, SMS*, and *PIN4*) and only 1/28 genes exhibit evidence for escaping (*CXorf36*). This means the other 24/28 show female-biased expression but no evidence of escape in the placenta.

In conclusion, we confirmed that XCI in the human placenta is skewed, but patchy, in contrast to adult tissues, which show mosaic XCI. In addition, we identified a subset of genes showing patterns of XCI that are unique to the placenta that should be further investigated for their roles in pregnancy and placentation.

## Materials and Methods

### Samples collection

Working with the Yale Biobank, we collected samples from 30 placentas, where the sex of the offspring was assigned female at birth. We confirmed that the samples were genetically XX by the presence of heterozygous sites across the X chromosome. The placenta samples were collected immediately following birth from term (≥ 36 weeks and 6 days) uncomplicated pregnancies delivered by cesarean section. Placentas were collected and sequenced at two different times, with 12 placenta samples in the first batch (OBG0044, OBG0068, OBG0111, OBG0115, OBG0120, OBG0133, OBG0156, OBG0170, OBG0174, OBG0175, OBG0178, and OBG0166) and 18 in the second (OBG0022, OBG0024, OBG0026, OBG0028, OBG0030, OBG0039, OBG0050, OBG0051, OBG0066, OBG0121, OBG0138, OBG0180, OBG0188, OBG0201, OBG0205, OBG0289, OBG0338, and OBG0342).

Placental tissue samples were obtained through rapid sampling (≤30 minutes post-delivery). Tissue samples were taken from the “maternal” side, midway between the chorionic and basal plates, from the periphery of the lobules, avoiding maternal tissue, consistent with best practices (Konwar et al. 2019) . Sampling sites were free of visible infarction, calcification, hematoma, and tears (areas of frank visible pathology were avoided). Whenever possible, tissue samples were obtained from distinct cotyledons of the placenta in opposing quadrants of the placenta (far from one another spatially).

The sampling protocol was as follows:

1. Orient the placenta maternal side up (basal plate uppermost) and identify sampling areas in each of the four placental quadrants.
2. At each site, remove the basal plate (to remove maternal tissue) ∼1-2mm by trimming with a pair of sterile scissors to expose villous tissue.
3. Cut a “grape-sized” (approximately 1-2 cm3; 5-6g) tissue lobule from each of the four quadrants.
4. Wash the tissue thoroughly twice in phosphate buffered saline (PBS) solution and blot on clean gauze.
5. From each “grape-sized” lobule, cut away eight smaller pieces (∼1-2 mm3) using a scalpel.
6. For each sampling quadrant, place four of the ∼50mg tissue pieces in a labeled cryovial. Snap freeze immediately in liquid nitrogen. Store samples at −80°C until ready for use; aliquot individual tissue pieces as needed per research protocols.
7. For each sampling quadrant, place four of the tissue pieces in a labeled cryovial containing 1 mL of RNAlater® solution (RNA stabilizing agent). Store the cryovials in the 4°C benchtop fridge for a minimum of 48 hours, and a maximum of 7 days, per RNAlater® manufacturer protocol. After a minimum of 48 hours, use a pipette to remove the RNAlater solution from the cryovials and immediately snap freeze in liquid nitrogen. Store samples at –80°C until ready for use; aliquot individual tissue pieces as needed per research protocols.

### Whole exome sequencing

DNA was extracted from one flash frozen collection site for each individual. Exome libraries were prepped, and sequenced to approximately 50X coverage with 100bp paired-end sequence on the Illumina NextSeq at the Yale Genome Sequence Center.

### Whole transcriptome sequencing

From each placenta, two separate sites from opposite quadrants were collected. RNA was extracted and RiboZero stranded libraries (RF) were prepared and sequenced to approximately 40 million reads per sample with 100bp paired-end sequence on the Illumina NovaSeq at the Yale Genome Sequence Center.

### Exome sequence data processing

We used fastqc version 0.11.8 (Andrews 2010) for quality control and aggregated results from fastqc by using multiqc version 0.9 (Ewels et al. 2016). We trimmed adapters using bbduk as part of bbmap version 38.22 (Bushnell 2014). Sequences were trimmed both left and right for phred quality of 30, minimum length of 75 base pairs, and average read quality of 20 or greater to keep only high quality reads (Bushnell 2014). Quality was checked after trimming (**Figure S1)**. We confirmed genetic XX females by examining the reads mapped ratio between the X chromosome and chromosome 19 (chrX/chr19), between the Y chromosome and chromosome 19 (chrY/chr19), and between the Y chromosome and the X chromosome (chrX/chrY). We observed that sample OBG0175 has a lower chrX/chr19 reads mapped ratio than other samples in this study (**Figure S2**). Therefore, we removed sample OBG0175 from further analyses. We used bwa-mem version 0.7.17 (Li 2013) to map to a female-specific reference genome. Specifically, we mapped the exome samples to a sex chromosome complement informed reference genome in which the Y chromosome is hard-masked (to avoid mismapping of X-linked reads to homologous regions on the Y chromosome in the XX samples). Mapping exome samples to a reference genome with the Y chromosome hard-masked has been shown to increase the number of variants genotyped on the X chromosome (Webster et al. 2019). To generate the sex chromosome complement reference genome we employed XYalign (Webster et al. 2019). XYalign created a Y-masked gencode GRCh38.p12 human reference genome for aligning XX individuals (Harrow et al. 2012). We used picard version 2.18.27 (“Picard Tools − By Broad Institute” n.d.) to mark PCR duplicates. To genotype variants, we used GATK version 4.1.0.0 (McKenna et al. 2010; DePristo et al. 2011; Van der Auwera et al. 2013). We first used GATK’s HaplotypeCaller to generate GVCF files. Second, we combined GVCF from 66 samples using GATK’s CombineGVCFs (30 placenta samples and an additional 36 samples from a separate study in the group (unpublished) here to increase power to genotype variants). Finally, we used GATK’s GenotypeGVCFs to call variants. Following GATK’s best practice, we obtained high-quality variants by filtering using GATK’s Variant Quality Score Recalibration (VQSR). We tabulated the number of heterozygous variants for each sample in **Table S1**.

Because we sequenced the placenta samples in two separate batches, we plotted the principal component for the exome data using the package *SNPRelate* in R (Zheng et al. 2012) and observed no separation by batch from the exome data (**Figure S3**).

### Whole transcriptome data processing

Samples were checked for quality using fastqc version 0.11.8 (Andrews 2010) and results were aggregated across samples using multiqc version 0.9 (Ewels et al. 2016). Adapters were removed and sequences were trimmed both left and right for phred quality of 30, minimum length of 75 base pairs, and average read quality of 20 or greater using bbduck version 38.22 (Bushnell 2014). Quality was checked after trimming (**Figure S4)**. Trimmed RNA-seq female placenta reads were then aligned to a sex chromosome complement informed reference genome with the Y chromosome hard masked with Ns (Webster et al. 2019; Olney et al. 2019) using the gencode GRCh38.p12 reference genome (Harrow et al. 2012). RNA-seq samples were aligned using HISAT2 (Kim, Langmead, and Salzberg, n.d.); which has been shown to be a robust aligner for high throughput RNA-seq reads. Total reads mapped and duplicate reads were visually checked using BAMtools stats (Barnett et al. 2011) (**Table S2)**.

### Obtaining allele counts for allele specific expression analysis

We used GATK ASEReadCounter (version 4.1.0.0) to obtain allele counts (McKenna et al. 2010) with the following thresholds: min-depth-of-non-filtered-base = 1, min-mapping-quality = 10, and min-base-quality = 10. GATK ASEReadCounter automatically filters out duplicated reads. Duplicated reads are suggested to be removed for allele specific expression analysis (Castel et al. 2015). The number of heterozygous and expressed variants for each sample are reported in **Table S3**.

### Genotype-Tissue Expression (GTEx) data processing

We downloaded ASEReadCounter counts for the X chromosome and chromosome 8 from the Genotype-Tissue Expression (GTEx) project, approved for project #8834 for General Research Use in Genotype-Tissue Expression (GTEx) to MAW. We only considered tissues with more than ten samples per tissue and excluded two non-primary tissues: EBV-transformed lymphocytes and cultured fibroblasts, leaving 45 tissues to be analyzed in this study. We list the GTEx tissues, the number of samples per tissue examined in this study, and the number of skewed samples per tissue in **Table S4**.

### Computing unphased median allele balance

For each heterozygous and expressed variant, ASEReadCounter tabulates the number of reads (the count) for the reference allele and the alternate allele. The total read count is the sum of the read count of the reference allele and the read count of the alternate allele. We define the biased allele to be the allele with more read counts. For each heterozygous and expressed variant, the unphased allele balance is the ratio between the read count of the biased allele and the total read count. Then, for each sample, the median allele balance is calculated by computing the median unphased allele balance across all heterozygous and expressed variants.

### Defining threshold for allele specific expression

To determine the threshold for defining biased allele expression, we plotted the unphased allele balance across all variants for each individual for chromosome 8 and the X chromosome (**Figure S5**). We observed that the unphased allele balance of most variants on the X chromosome is greater than 0.8 while the unphased allele balance of most variants on chromosome 8 is less than 0.8 (**Figure S5**). Therefore, in this study, we used 0.8 as a threshold for biased allele specific expression.

### Determining which X chromosome is inactivated

To determine whether the same X chromosome is inactivated at the two extraction sites for each placenta sample, we employed a phasing strategy on the X chromosome by defining that the biased alleles (the alleles with higher counts) are on the same haplotype. We restricted this analysis to contain only heterozygous and expressed variants that are shared between the two extraction sites. For consistency, we defined extraction site A to be the site with more expressed variants where the unphased allele balance is greater than 0.8. We defined the activated X chromosome to consist of alleles where its allele balance (the ratio between this allele’s read count and the total read count) is greater than 0.8. In cases where the allele balance is less than 0.8, we picked an allele at random with equal probability between the reference allele and the alternate allele. Then, we computed the allele balance (defined as the ratio between the biased allele’s count and total count) for the alleles on the same X chromosome. We called this phased allele balance. At each site, we computed the median phased allele balance to use as summary statistics. To compute phased allele balance we followed this procedure:

1. Find expressed variants that are shared between site A and site B.
2. Using the shared expressed variants between site A and site B, tabulate the number of expressed variants that exhibit biased expression (i.e., unphased allele balance is greater than 0.8).
3. Pick a site to base phasing from based on the number of expressed variants with biased expression. For example, if site A has 50 expressed variants with biased expression and site B has 60 such variants, phasing is based on site B.
4. Phasing strategy (using site B to base phasing): the expressed haplotype is generated by calculate: For each expressed variant that is shared between site A and site B, pick the allele with allele balance greater than 0.8 to be on the expressed haplotype. If allele balance is less than 0.8, choose an allele at random with equal probability

### Validating method based on allele specific expression to quantify XCI

We validated our approach to determine skewness by examining the non-pseudoautosomal regions (nonPARs) of the X chromosome in males. We collected samples from 12 placentas, where the sex of the offspring was assigned male at birth. Even though the X chromosome in males is haploid, some variants are incorrectly genotyped as being heterozygous when diploidy is assumed when genotyping with GATK (**Table S5**). Regardless, these heterozygous variants exhibit mostly skewed expression, indicating that our method of combining both the whole exome and whole transcriptome sequence data is robust to determine skewness (**Figure S6**).

### Classifying genes into genes that are inactivated, genes that escape XCI, and genes that show variable escape

For classifying genes, we only considered samples that show skewed expression (allele balance is greater than 0.8). There are 52/58 samples in the placenta dataset and 525/4958 samples in the GTEx dataset exhibiting skewed expression that were used for this analysis. For each gene on the X chromosome, we only considered a gene if there are at least five informative samples where in each sample, there is at least one heterozygous and expressed variant for that gene. If there are more than one heterozygous and expressed variant for a gene, we summed up the counts for the biased allele and calculated allele balance by dividing the summed counts of the biased allele by the summed counts of the total counts. We categorized a gene as being inactivated if its allele balance is greater or equal to 0.8 in at least 70% of the samples and the median allele balance across all samples is greater or equal to 0.8. We categorized a gene as escaping from XCI if at most 30% of the samples have allele balance for this gene equal or greater than 0.8 and the median allele balance across all samples is less than or equal to 0.75. Otherwise, the gene is annotated as showing variable escape across all individuals. See **Supplementary Note 2** for more information.

### Coordinates used of PARs and XIST

As defined for GRCh38.p12 (“Human Genome Overview - Genome Reference Consortium” n.d.), we used the following coordinates for the pseudoautosomal region 1 (PAR1), PAR2, and XIST.

PAR1: position 10,001 to position 2,781,479 on the X chromosome

PAR2: position 155,701,383 to position 156,030,895 on the X chromosome

XIST: position 73,820,651 to position 73,852,723 on the X chromosome

## Acknowledgements

We acknowledge Research Computing at Arizona State University for providing high-performance computing and storage resources that have contributed to the research results reported within this paper. URL: http://www.researchcomputing.asu.edu. We would also like to thank Heini Natri and Wilson lab members for helpful feedback on the manuscript. This work was supported by the National Institute of General Medical Sciences (NIGMS) of the National Institutes of Health (NIH) grant R35GM124827 to MAW. This research was supported by The Yale University Reproductive Sciences Data and Specimen Biorepository, HIC#1309012696, a component of the Department of Obstetrics, Gynecology & Reproductive Sciences, Yale School of Medicine, New Haven, CT. Research reported in this publication was supported by the Eunice Kennedy Shriver National Institute Of Child Health & Human Development of the National Institutes of Health under Award Number F31HD101252 to KCO. KCO was additionally supported by ARCS Spetzler Scholar.

## Data and Code Availability

The datasets generated during this study are available via controlled access: phs002240.v1.p1 NIGMS Sex Differences Placentas. The code generated during this study is available at https://github.com/SexChrLab/Placenta_XCI.

