## Supplemental Material for "X chromosome inactivation in the human placenta is patchy and distinct from adult tissues"

### Title

### Authors and Affiliations

Tanya N. Phung[1,2], Kimberly C. Olney[1,2], Michelle Silasi[3], Lauren Perley[4], Jane O'Bryan[4], Harvey J. Kliman[4], and Melissa A. Wilson[1,2,5]

1. Center for Evolution and Medicine, Arizona State University, Tempe AZ 85282 USA
2. School of Life Sciences, Arizona State University, Tempe AZ 85282 USA
3. Department of Maternal-Fetal Medicine, Mercy Hospital St. Louis, St. Louis, MO 63141
4. Department of Obstetrics, Gynecology and Reproductive Sciences, Yale University School of Medicine, New Haven, CT 06520
5. The Biodesign Center for Mechanisms of Evolution, Arizona State University, Tempe AZ 85282 USA

### Corresponding Author

Melissa A. Wilson
School of Life Sciences | Arizona State University | PO Box 874501 | Tempe, AZ 85287-4501


### Supplementary Information

#### Table S1. Number of heterozygous variants on the X chromosome and chromosome 8

For each sample, we reported the number of heterozygous variants after variant genotyping using GATK and after filtering using VQSR.

| **SampleID** | **X chromosome** | **Chromosome 8** |
|---|---|---|
| OBG0044 | 951 | 1691 |
| OBG0068 | 1468 | 2218 |
| OBG0111 | 832 | 1650 |
| OBG0115 | 964 | 1568 |
| OBG0120 | 824 | 1674 |
| OBG0133 | 1316 | 2134 |
| OBG0156 | 997 | 1600 |
| OBG0170 | 1390 | 1979 |
| OBG0174 | 1426 | 2158 |
| OBG0175 | 607 | 1594 |
| OBG0178 | 1157 | 1906 |
| OBG0166 | 804 | 1620 |
| OBG0022 | 1125 | 1964 |
| OBG0024 | 1206 | 2152 |
| OBG0026 | 1907 | 2912 |
| OBG0028 | 1706 | 2730 |
| OBG0030 | 1842 | 3126 |
| OBG0039 | 1086 | 2359 |
| OBG0050 | 1786 | 2553 |
| OBG0051 | 1107 | 2154 |
| OBG0066 | 2010 | 2906 |
| OBG0121 | 1805 | 2747 |
| OBG0138 | 1044 | 2268 |
| OBG0180 | 1110 | 2257 |
| OBG0188 | 1940 | 2874 |
| OBG0201 | 1757 | 2741 |
| OBG0205 | 1275 | 2393 |
| OBG0289 | 2020 | 2805 |
| OBG0338 | 1049 | 2093 |
| OBG0342 | 1169 | 2139 |

#### Table S2. Mapped reads at each extraction site for the whole transcriptome and for the X chromosome.

For each sample, we used samtools stats to obtain the number of reads that mapped.

| Sample ID | Site A | | Site B | |
|---|---|---|---|---|
| | **Whole transcriptome** | **Chr X** | **Whole transcriptome** | **Chr X** |
| OBG0044 | 124821361 | 2570404 | 104951941 | 1956346 |
| OBG0068 | 96468961 | 2028381 | 72630017 | 1632320 |
| OBG0111 | 105360653 | 2210820 | 87310785 | 1746585 |
| OBG0115 | 115047845 | 2233186 | 81406062 | 1794380 |
| OBG0120 | 109637796 | 2360408 | 78369660 | 1653373 |
| OBG0133 | 106542634 | 2158139 | 75940154 | 1709600 |
| OBG0156 | 123238340 | 2617278 | 79648267 | 1954931 |
| OBG0170 | 104502826 | 2539731 | 101090503 | 2263802 |
| OBG0174 | 29178948 | 589785 | 95550124 | 1787220 |
| OBG0175 | 98337411 | 2134510 | 85986263 | 2061137 |
| OBG0178 | 114938081 | 2447404 | 86675194 | 1916321 |
| OBG0166 | 114474479 | 2532057 | 93337543 | 2043997 |
| OBG0022 | 79624620 | 1846538 | 77386563 | 2048223 |
| OBG0024 | 56525905 | 1464827 | 136862205 | 3467075 |
| OBG0026 | 59648434 | 1014174 | 86201051 | 1772717 |
| OBG0028 | 69962192 | 1894168 | 42201549 | 964035 |
| OBG0030 | 75533477 | 1797735 | 72423761 | 1799896 |
| OBG0039 | 81923032 | 1986227 | 60580683 | 1631386 |
| OBG0050 | 72749515 | 1711200 | 62546230 | 1564591 |
| OBG0051 | 78292464 | 1885179 | 61467811 | 1513737 |
| OBG0066 | 74368314 | 1535224 | 80264229 | 2054109 |
| OBG0121 | 95299259 | 2477900 | 40005508 | 1132595 |
| OBG0138 | 217429809 | 5692793 | 73761980 | 1578129 |
| OBG0180 | 72664680 | 1974885 | 29697974 | 837063 |
| OBG0188 | 21675595 | 617351 | 79407643 | 2295587 |
| OBG0201 | 88301001 | 2190583 | 80598634 | 2085968 |
| OBG0205 | 49073152 | 1316484 | 40796256 | 1070053 |
| OBG0289 | 72176230 | 1850324 | 78471721 | 1954236 |
| OBG0338 | 75400996 | 1972344 | 57209598 | 1381726 |
| OBG0342 | 80308862 | 1916568 | 85785465 | 2074156 |

#### Table S3. Number of heterozygous and expressed variants on the non-pseudoautosomal regions of the X chromosome and chromosome 8

After running GATK ASEReadCounter, we tabulated the number of heterozygous variants that are expressed (where total RNA read count is greater than 10).

| **Sample ID** | **Chromosome X** | | **Chromosome 8** | |
|---|---|---|---|---|
| | **Site A** | **Site B** | **Site A** | **Site B** |
| OBG0044 | 97 | 87 | 219 | 214 |
| OBG0068 | 100 | 94 | 251 | 258 |
| OBG0111 | 74 | 65 | 170 | 161 |
| OBG0115 | 76 | 68 | 166 | 151 |
| OBG0120 | 74 | 57 | 241 | 197 |
| OBG0133 | 103 | 94 | 239 | 211 |
| OBG0156 | 100 | 93 | 218 | 222 |
| OBG0170 | 120 | 124 | 270 | 268 |
| OBG0174 | 62 | 120 | 116 | 239 |
| OBG0175 | 61 | 87 | 167 | 182 |
| OBG0178 | 89 | 87 | 223 | 219 |
| OBG0166 | 89 | 67 | 193 | 176 |
| OBG0022 | 119 | 112 | 234 | 244 |
| OBG0024 | 100 | 212 | 177 | 344 |
| OBG0026 | 109 | 125 | 262 | 273 |
| OBG0028 | 92 | 159 | 153 | 253 |
| OBG0030 | 165 | 168 | 362 | 373 |
| OBG0039 | 68 | 94 | 225 | 297 |
| OBG0050 | 160 | 148 | 256 | 274 |
| OBG0051 | 100 | 91 | 249 | 214 |
| OBG0066 | 160 | 127 | 422 | 355 |
| OBG0121 | 186 | 95 | 338 | 176 |
| OBG0138 | 191 | 77 | 534 | 212 |
| OBG0180 | 92 | 43 | 262 | 133 |
| OBG0188 | 59 | 183 | 111 | 336 |
| OBG0201 | 125 | 131 | 304 | 328 |
| OBG0205 | 71 | 61 | 169 | 144 |
| OBG0289 | 161 | 171 | 319 | 316 |
| OBG0338 | 87 | 63 | 244 | 173 |
| OBG0342 | 92 | 96 | 255 | 288 |

#### Table S4. Number of samples for each adult GTEx tissue.

The number of samples for each adult GTEx tissue used in this study (column 2), and the number of skewed samples (median allele balance greater than 0.8) (column 3).

| **GTEx tissues** | **Number of samples** | **Number of skewed samples** |
|---|---|---|
| Adipose_Subcutaneous | 194 | 19 |
| Adipose_Visceral_Omentum | 149 | 11 |
| Adrenal_Gland | 94 | 18 |
| Artery_Aorta | 138 | 7 |
| Artery_Coronary | 84 | 11 |
| Artery_Tibial | 187 | 14 |
| Brain_Amygdala | 37 | 2 |
| Brain_Anterior_cingulate_cortex_BA24 | 42 | 2 |
| Brain_Caudate_basal_ganglia | 52 | 3 |
| Brain_Cerebellar_Hemisphere | 51 | 5 |
| Brain_Cerebellum | 58 | 4 |
| Brain_Cortex | 64 | 4 |
| Brain_Frontal_Cortex_BA9 | 48 | 2 |
| Brain_Hippocampus | 49 | 2 |
| Brain_Hypothalamus | 47 | 1 |
| Brain_Nucleus_accumbens_basal_ganglia | 55 | 4 |
| Brain_Putamen_basal_ganglia | 42 | 3 |
| Brain_Spinal_cord_cervical_c-1 | 48 | 1 |
| Brain_Substantia_nigra | 33 | 2 |
| Breast_Mammary_Tissue | 151 | 20 |
| Colon_Sigmoid | 113 | 3 |
| Colon_Transverse | 136 | 15 |
| Esophagus_Gastroesophageal_Junction | 110 | 3 |
| Esophagus_Mucosa | 176 | 32 |
| Esophagus_Muscularis | 162 | 9 |
| Heart_Atrial_Appendage | 119 | 9 |
| Heart_Left_Ventricle | 122 | 21 |
| Kidney_Cortex | 18 | 0 |
| Liver | 62 | 12 |
| Lung | 166 | 13 |
| Minor_Salivary_Gland | 40 | 4 |
| Muscle_Skeletal | 237 | 17 |
| Nerve_Tibial | 177 | 13 |

| | | |
|---|---|---|
| Ovary | 167 | 21 |
| Pancreas | 116 | 23 |
| Pituitary | 71 | 4 |
| Skin_Not_Sun_Exposed_Suprapubic | 169 | 28 |
| Skin_Sun_Exposed_Lower_leg | 208 | 35 |
| Small_Intestine_Terminal_Ileum | 63 | 5 |
| Spleen | 86 | 4 |
| Stomach | 122 | 15 |
| Thyroid | 196 | 14 |
| Uterus | 129 | 10 |
| Vagina | 141 | 20 |
| Whole_Blood | 229 | 60 |

Table S5. Number of heterozygous sites identified in nonPARs in XY males.

| Sample | Number of called variants (AC > 0) | Number of heterozygous variants (AC = 1) | Number of heterozygous & expressed variants in site A | Number of heterozygous & expressed variant in site B |
|---|---|---|---|---|
| OBG0112 | 1,479 | 213 | 27 | 32 |
| OBG0116 | 2,018 | 174 | 18 | 18 |
| OBG0117 | 1,343 | 149 | 31 | 27 |
| OBG0118 | 1,484 | 180 | 24 | 25 |
| OBG0122 | 1,413 | 126 | 15 | 11 |
| OBG0123 | 1,762 | 149 | 22 | 23 |
| OBG0126 | 1,898 | 171 | 20 | 20 |
| OBG0130 | 1,566 | 197 | 23 | 28 |
| OBG0132 | 1,946 | 182 | 20 | 18 |
| OBG0158 | 1,620 | 223 | 28 | 27 |
| YPOPS0006 | 1,510 | 180 | 26 | 27 |
| OBG0053 | 1,532 | 172 | 23 | 30 |

### Supplementary Figures

#### Figure S1. Whole exome post-trimming FastQC Mean Quality Scores.

We used fastQC to check for quality of reads after trimming and used multiQC to aggregate results (A) 12 placenta samples from batch 1. (B) 18 placenta samples from batch 2.

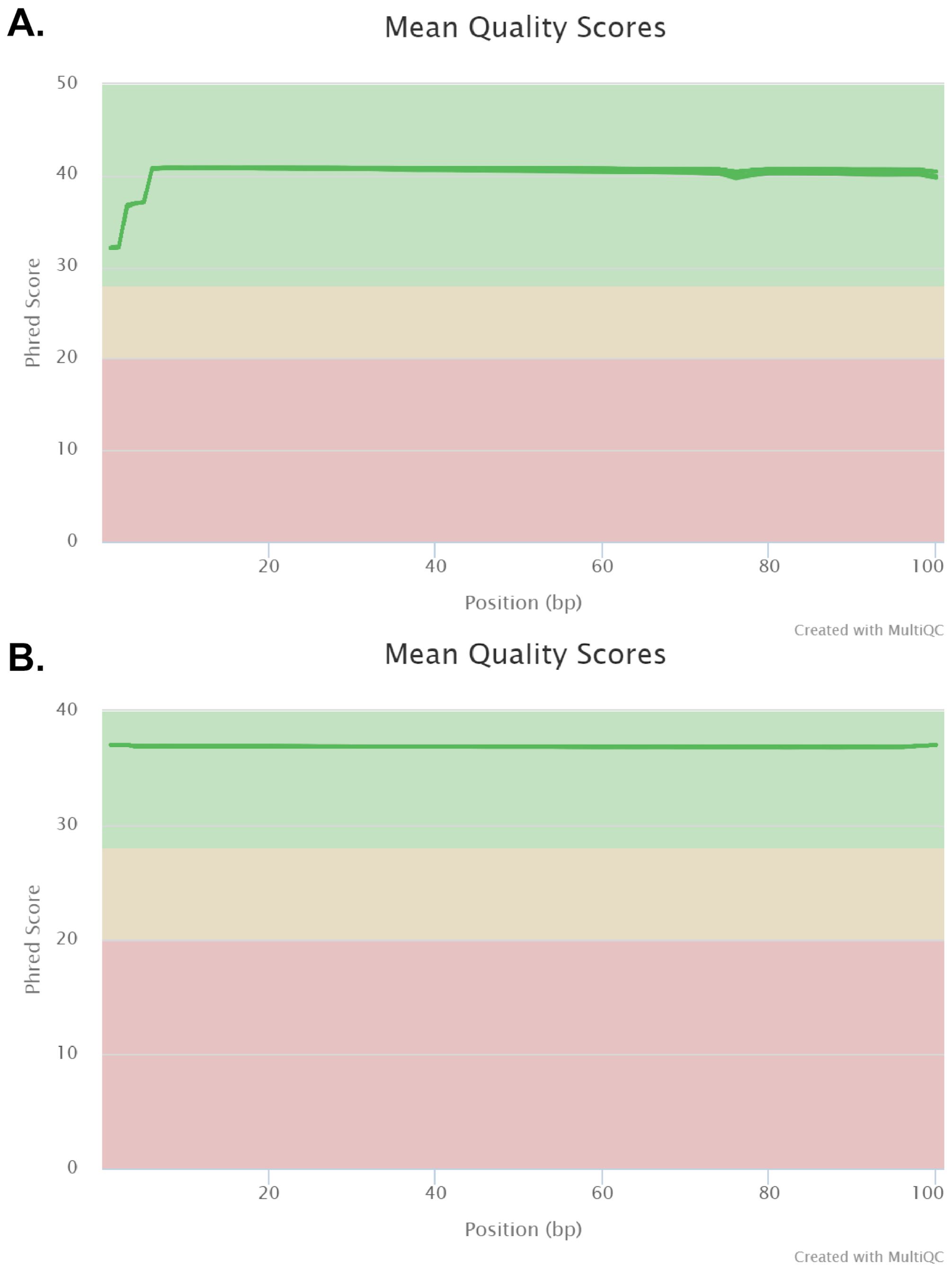

### Figure S2. Reads mapped ratio.

The number of reads mapped to the X chromosome, the Y chromosome, and chromosome 19 was obtained by running samtools stats. Ratios in reads mapped were plotted for between the X chromosome and chromosome 19 (chrX/chr19), between the Y chromosome and chromosome 19 (chrY/chr19), and between the Y chromosome and the X chromosome (chrY/chrX). We observed that the ratio of reads mapped ratio between the X chromosome and chromosome 19 is much lower for OBG0175 than all other samples, suggesting that this sample is not genetic XX sample. Therefore, we removed this sample from further analyses.

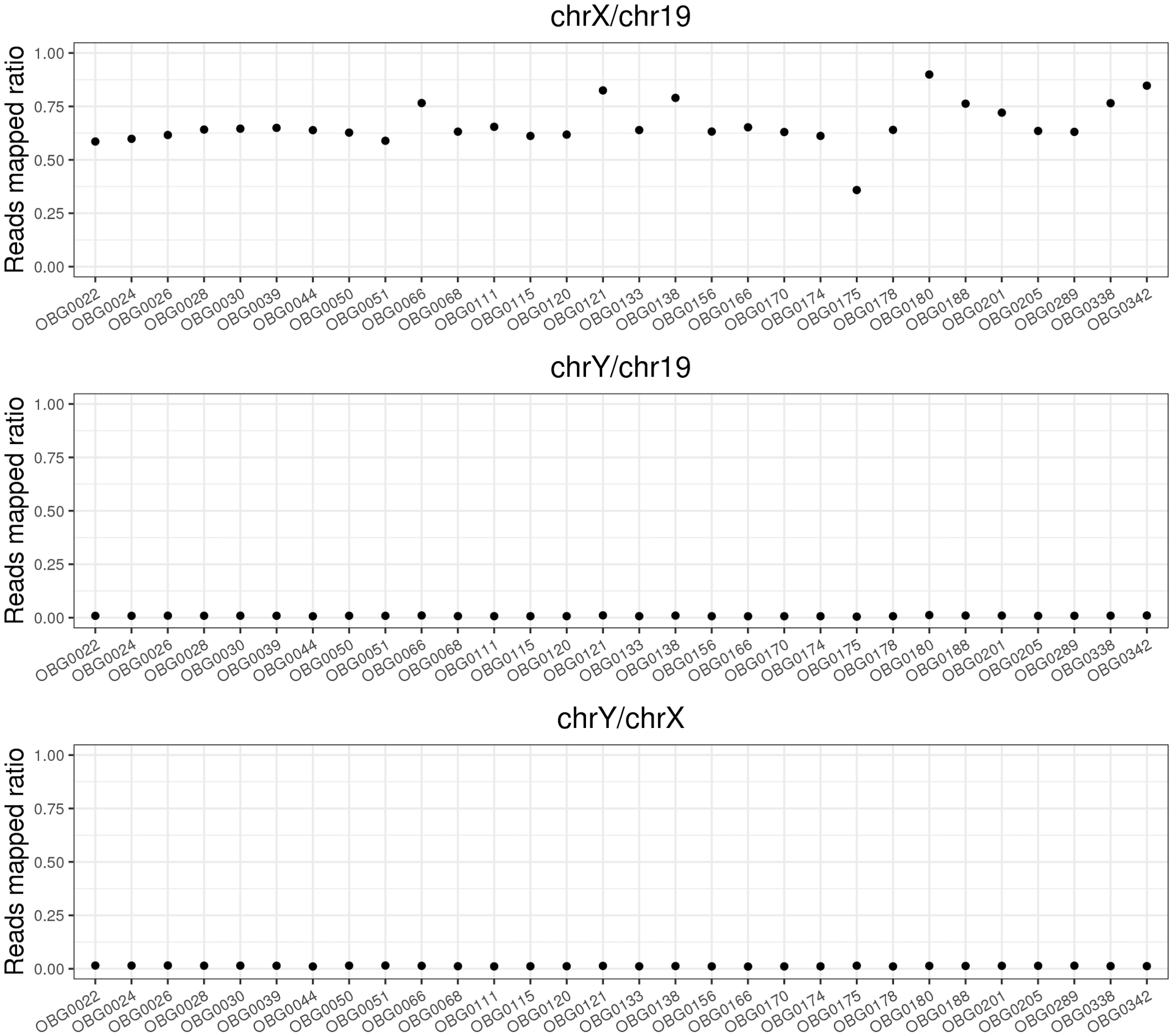

#### Figure S3. Principal component analysis for placenta samples.

Principal component for principal component 1 and 2 for the X chromosome (left panel) and for chromosome 8 (right panel) using the exome data from batch 1 and batch 2. We observed no clear separation by batches between these samples. The batch 1 samples are: OBG0044, OBG0068, OBG0111, OBG0115, OBG0120, OBG0133, OBG0156, OBG0170, OBG0174, OBG0175, OBG0178, and OBG0166. The batch 2 samples are: OBG0022, OBG0024, OBG0026, OBG0028, OBG0030, OBG0039, OBG0050, OBG0051, OBG0066, OBG0121, OBG0138, OBG0180, OBG0188, OBG0201, OBG0205, OBG0289, OBG0338, and OBG0342.

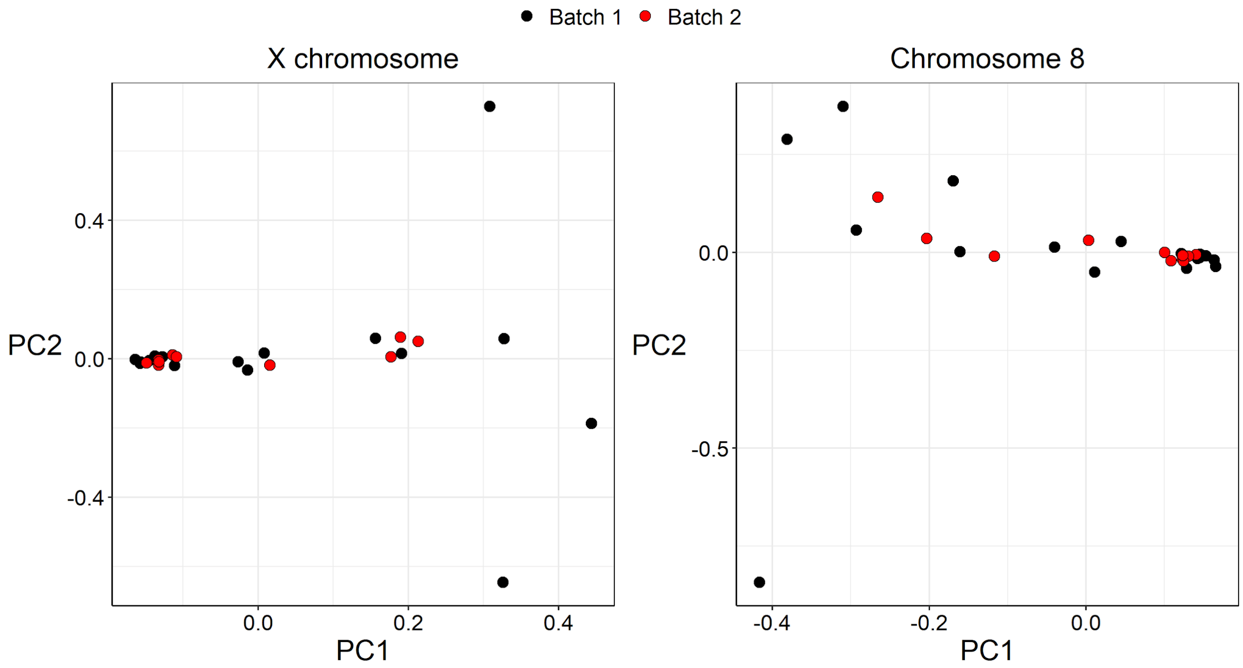

#### Figure S4. Whole transcriptome post-trimming FastQC Mean Quality Scores.

We used fastQC to check for quality of reads after trimming and used multiQC to aggregate results (A) 12 placenta samples from batch 1. (B) 18 placenta samples from batch 2.

**A.**

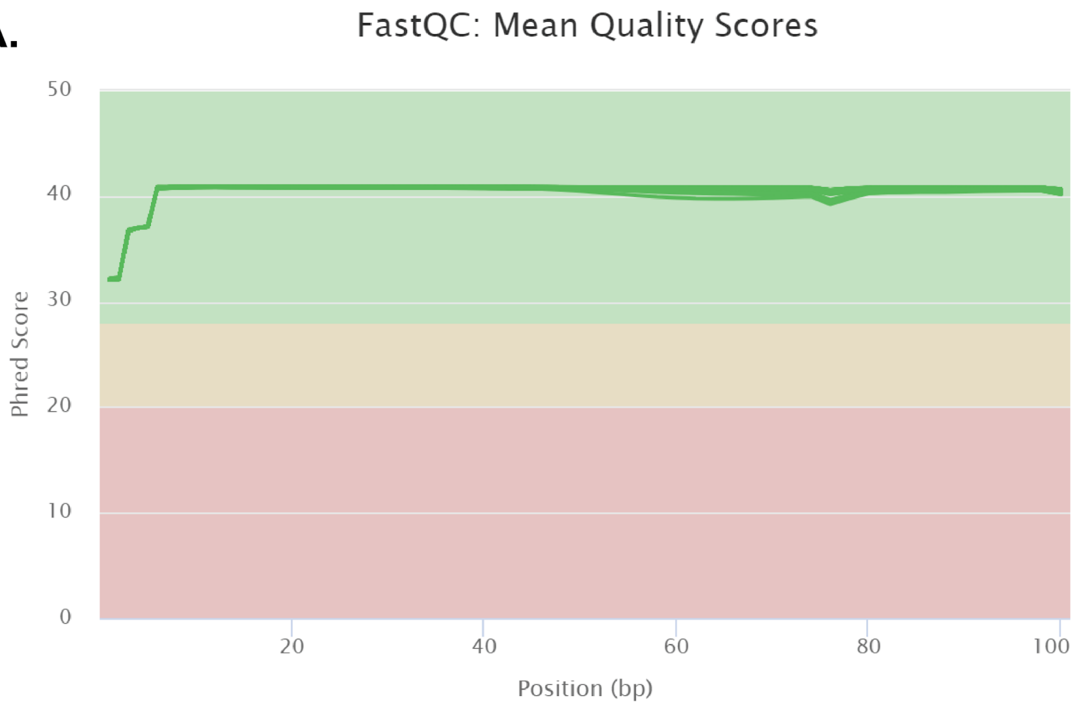


**B.**

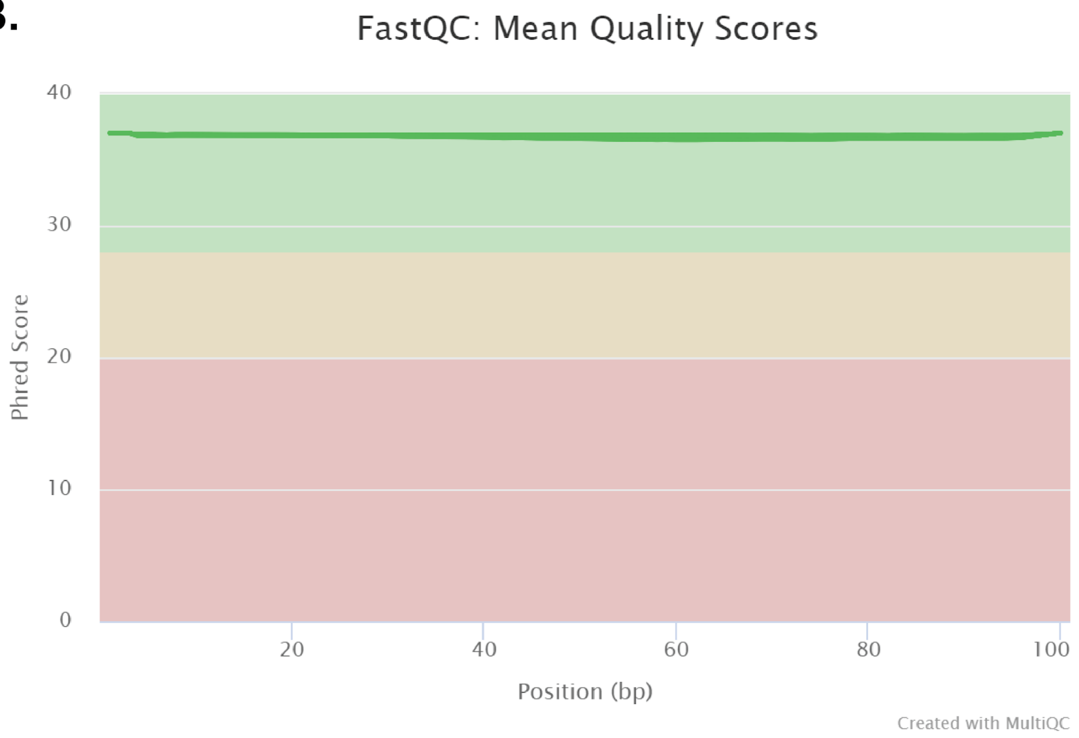

### Figure S5. Determining threshold for skewed inactivation by comparing allele balance between chromosome 8 and the X chromosome.

Each plot is a histogram of the allele balance for chromosome 8 (gray bars) and the X chromosome (yellow bars) for site A (left) and site B (right). Dotted lines denote allele balance of 0.8. We observed that the allele balance of most variants on chromosome 8 is less than 0.8 while the allele balance of most variants on the X chromosome is greater than 0.8.

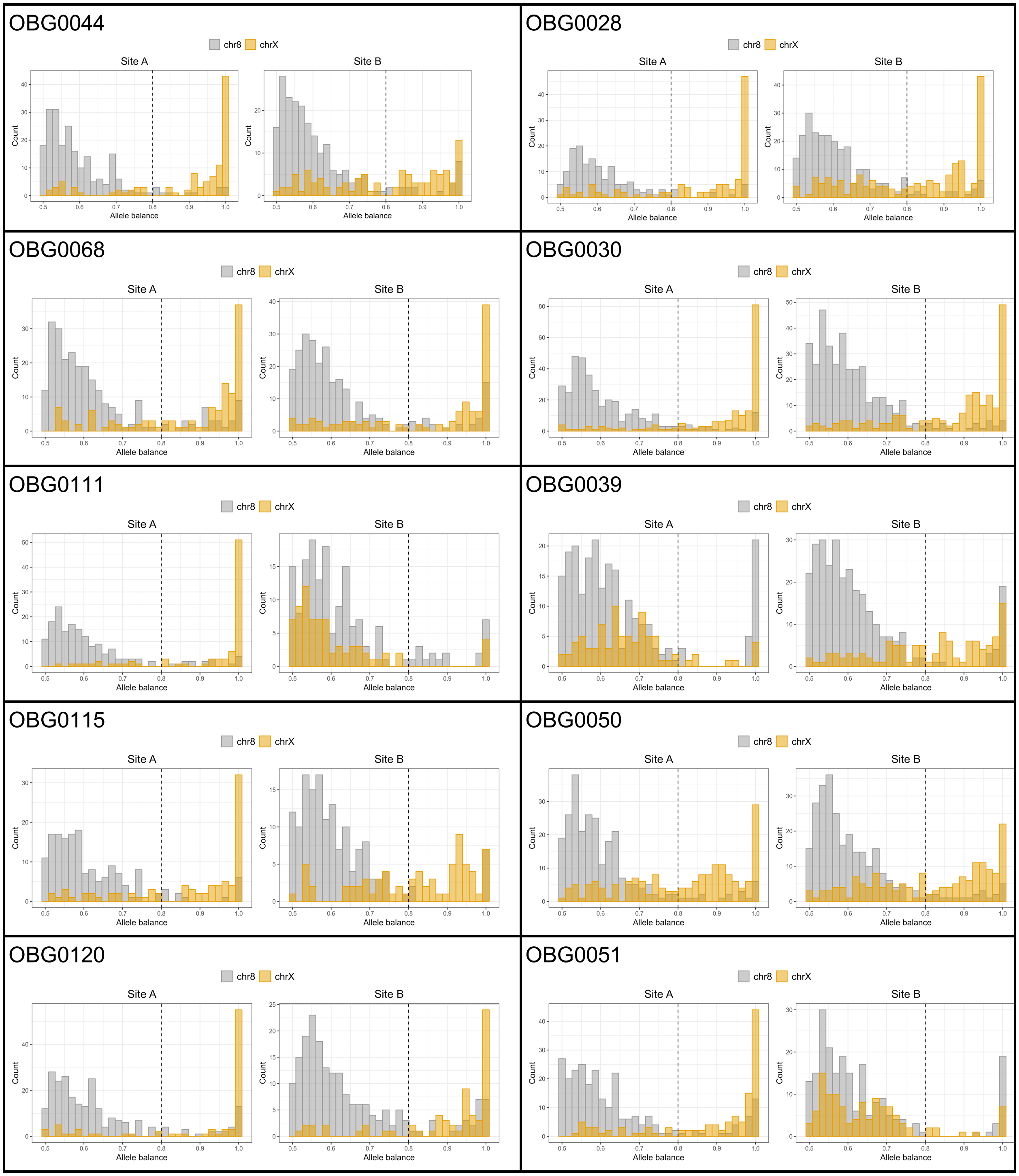

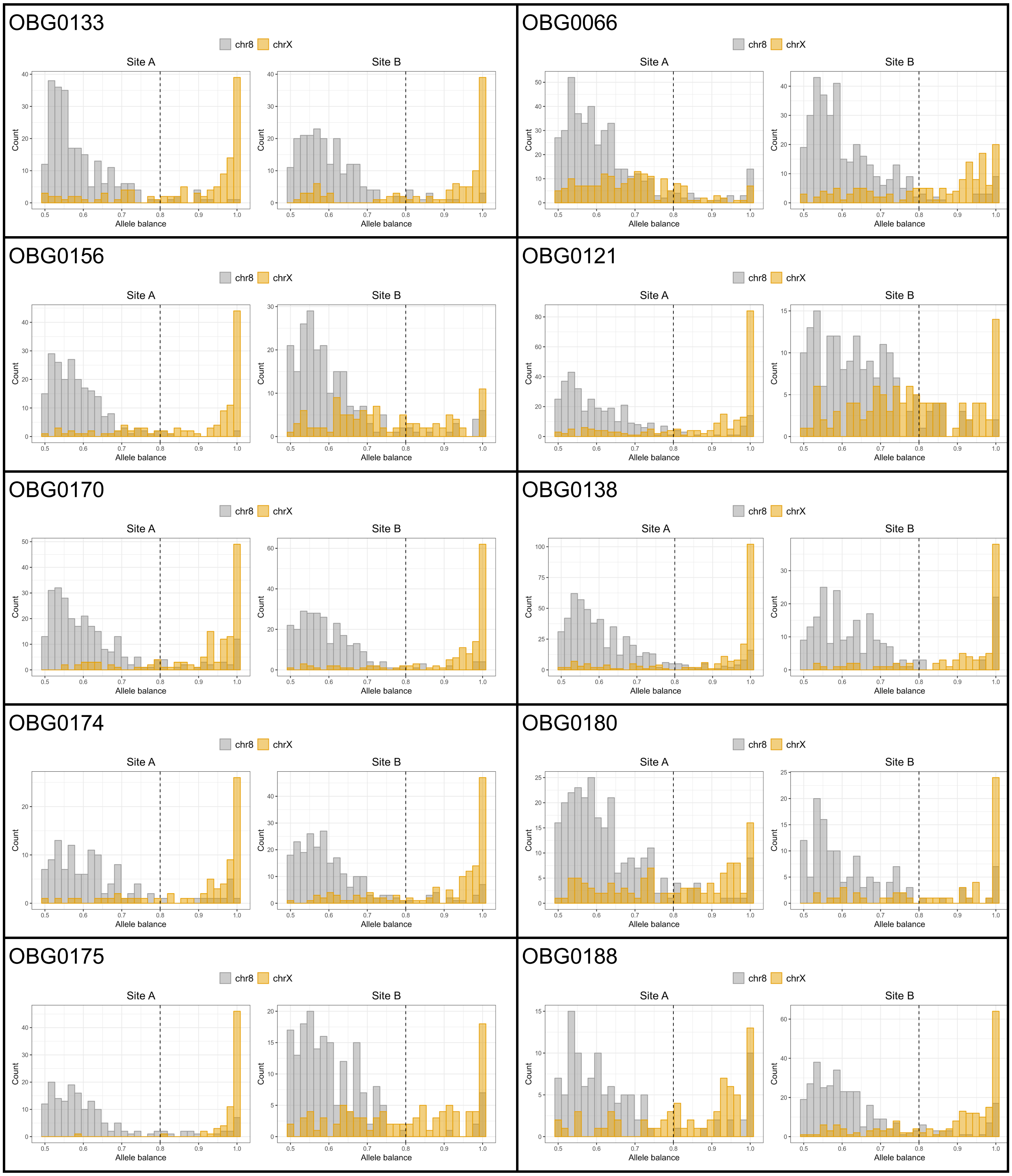
OBG0133
chr8
chrX
Site A
Site B
Count
Allele balance
OBG0066
OBG0156
OBG0121
OBG0170
OBG0138
OBG0174
OBG0180
OBG0175
OBG0188

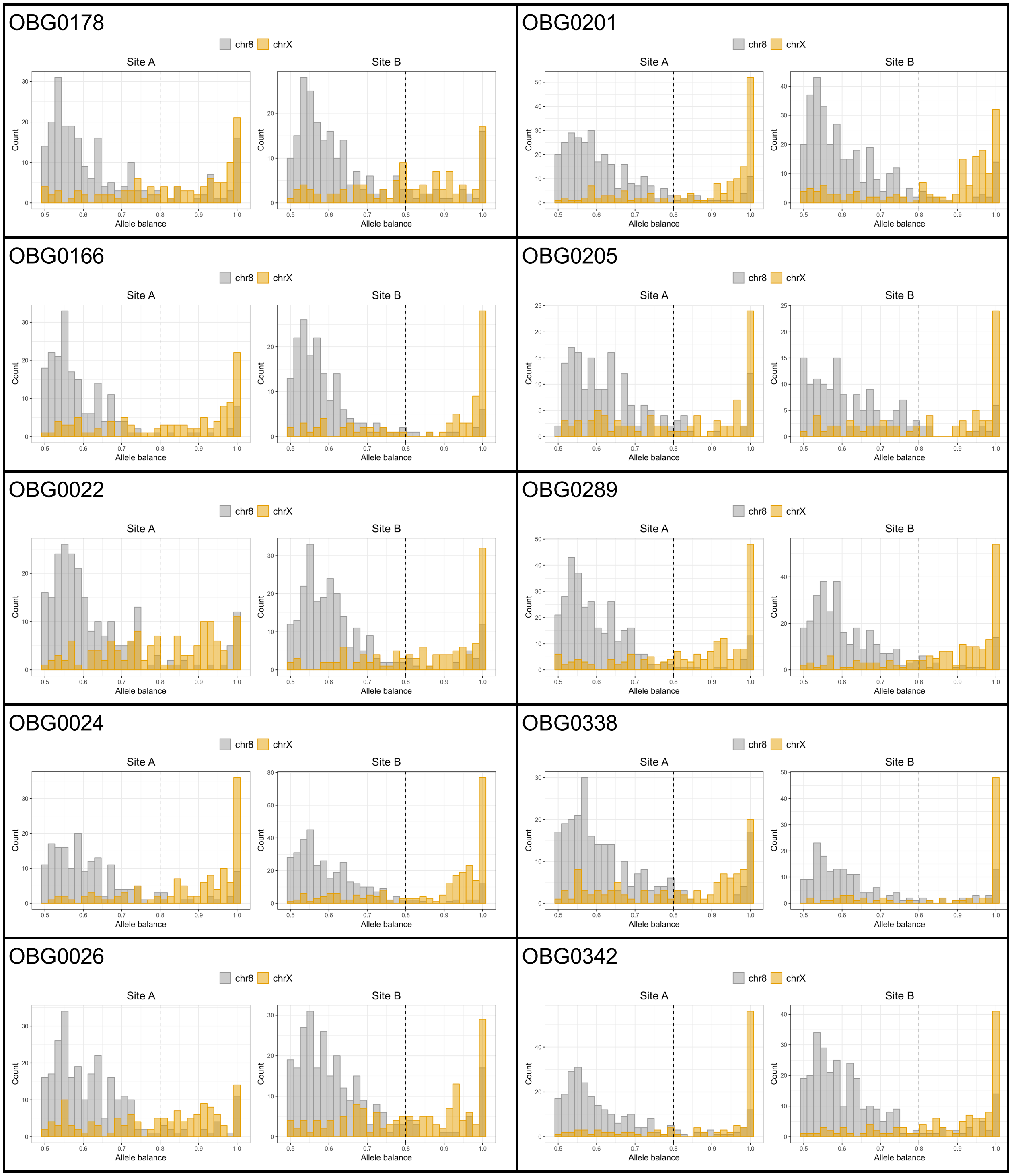
OBG0178
chr8
chrX
Site A
Site B
Count
Allele balance
OBG0201
chr8
chrX
Site A
Site B
Count
Allele balance
OBG0166
chr8
chrX
Site A
Site B
Count
Allele balance
OBG0205
chr8
chrX
Site A
Site B
Count
Allele balance
OBG0022
chr8
chrX
Site A
Site B
Count
Allele balance
OBG0289
chr8
chrX
Site A
Site B
Count
Allele balance
OBG0024
chr8
chrX
Site A
Site B
Count
Allele balance
OBG0338
chr8
chrX
Site A
Site B
Count
Allele balance
OBG0026
chr8
chrX
Site A
Site B
Count
Allele balance
OBG0342
chr8
chrX
Site A
Site B
Count
Allele balance

#### Figure S6. Most variants in the non-pseudoautosomal regions of the X chromosome in XY males are skewed.

Histogram of allele balance in nonPARs in male XY samples called as diploid. We joint-called genotypes on 12 male XY placentas (see **Methods**). The expression of variants on the nonPARs of the X chromosome in XY males should be completely biased towards only one allele because there is only one X chromosome. However, if we called the nonPARs as diploid, we wrongly identified only a small number of variants to be heterozygous (**Table S5**).

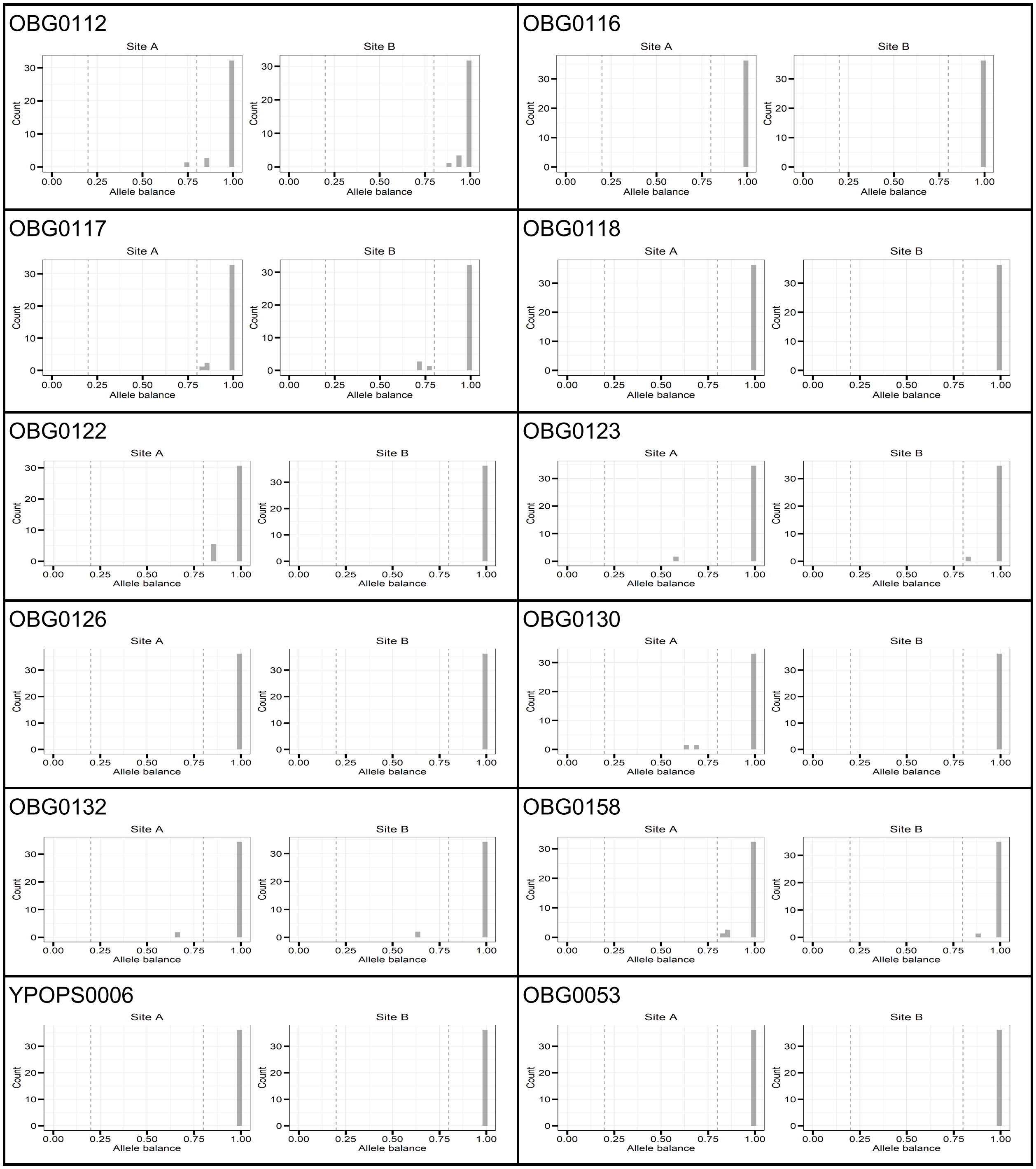

#### Figure S7. Patterns of X-inactivation across the entire X chromosome.

In each plot, allele balance at each heterozygous and expressed variant is plotted as a function of the position on the X chromosome. Open black circles denote variants on extraction site A. Filled red triangles denote variants on extraction site B. Gray boxes denote the pseudoautosomal regions and XIST.

A. Both extraction sites show the same X chromosome being inactivated

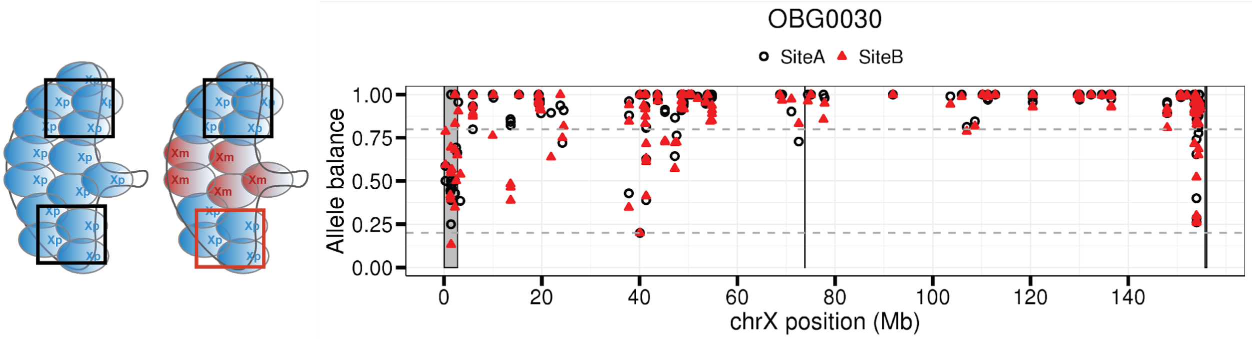


B. Each site shows a different X chromosome being inactivated

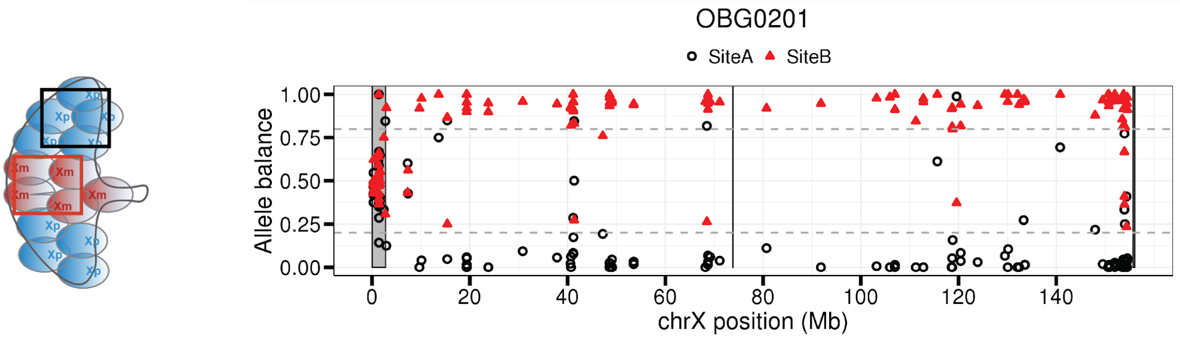


C. One extraction site shows skewed X-inactivation and the other shows both X chromosome being expressed

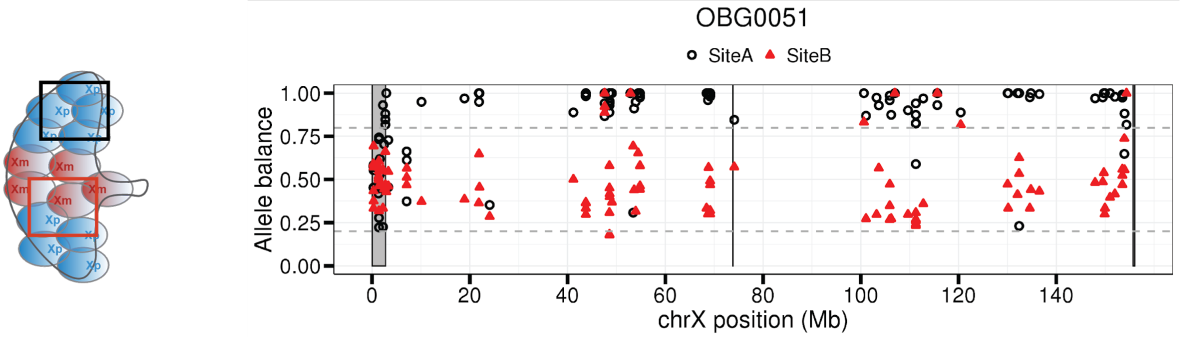

### Figure S8. Chromosome 8 shows biallelic expression in placenta and adult tissues.

Unphased median allele balance is plotted for the placenta in this study (purple) and for 45 adult tissues in the GTEx dataset (blue) on chromosome 8. Each point of the violin plot is the median allele balance for each sample. Because there are no multiple site samplings for the GTEx data, unphased allele balance was computed.

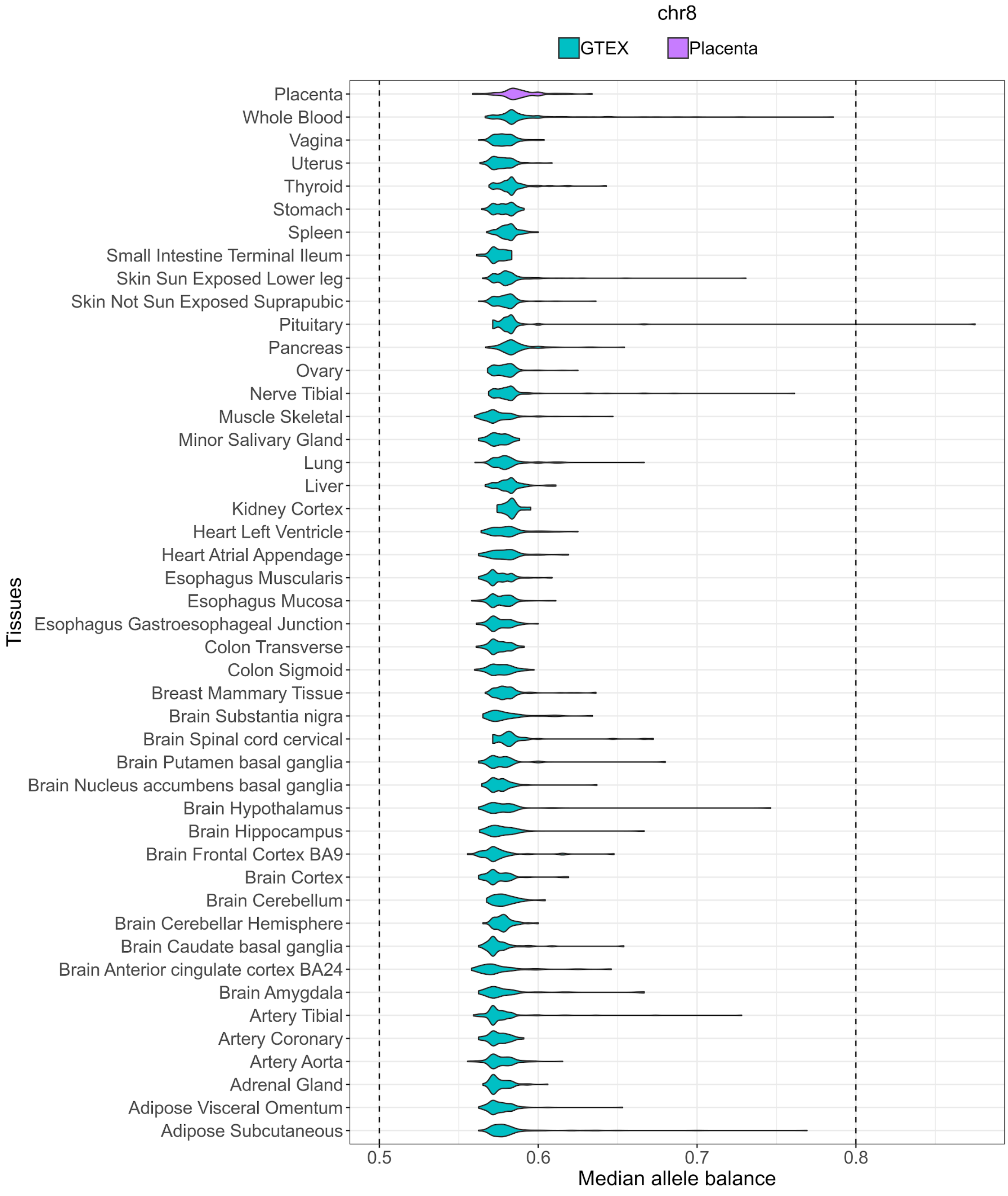

Figure S9. Heterogeneity in proportion of samples per gene that escape X-inactivation or are silenced.

For each gene, the dark blue bar denotes the proportion of samples that show evidence for that gene escaping XCI. The yellow bar denotes the proportion of samples that show evidence for that gene being silenced.

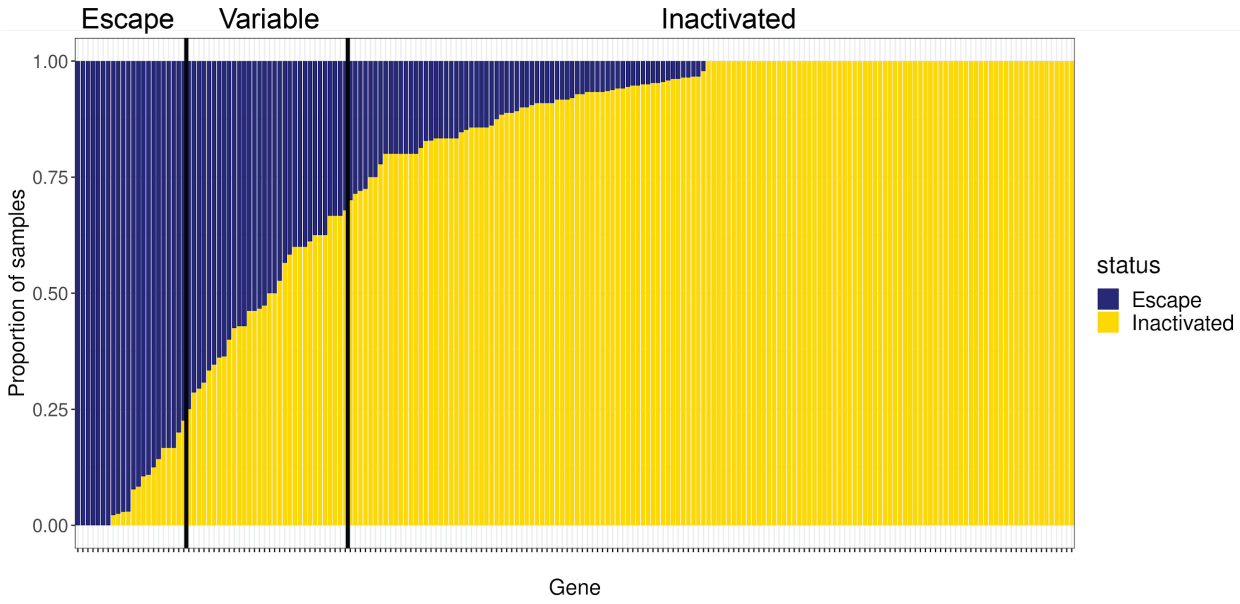

Figure S10. Heterogeneity in escape from X chromosome inactivation across and within placentas.

For each gene on the X chromosome, inactivation status is shown for each extraction site for each sample: dark blue (genes that escape XCI) and yellow (genes that are inactivated).

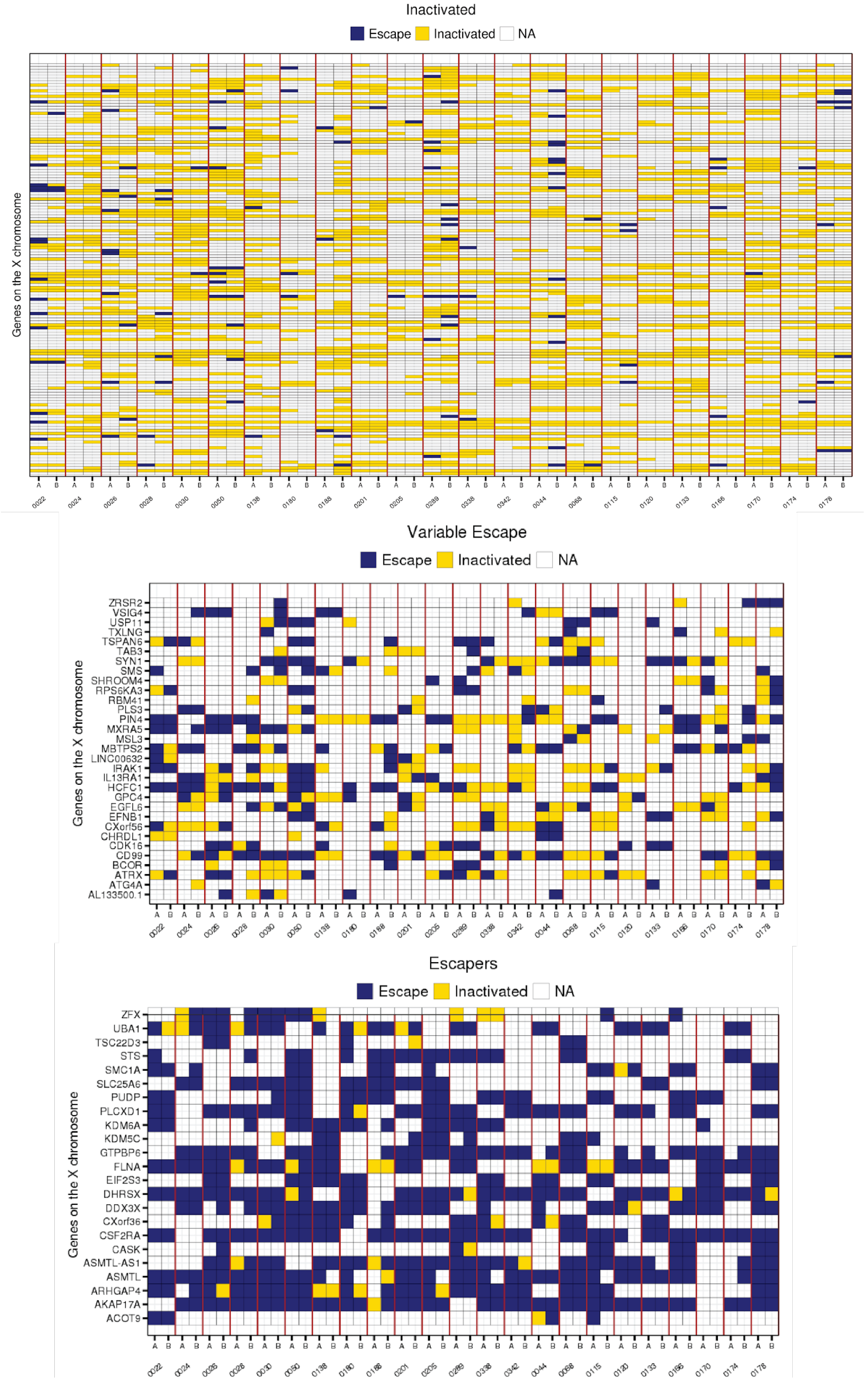

#### Figure S11. Higher gene expression in female does not necessarily equate to escape gene

Female to male $\log_2$ ratio (calculated as $\log_2(\text{female}_{CPM}/\text{male}_{CPM})$) in gene expression was computed for genes categorized as inactivated, escape, or variable in the placenta. The $\text{female}_{CPM}$ and $\text{male}_{CPM}$ were obtained from Olney et al. (unpublished data).

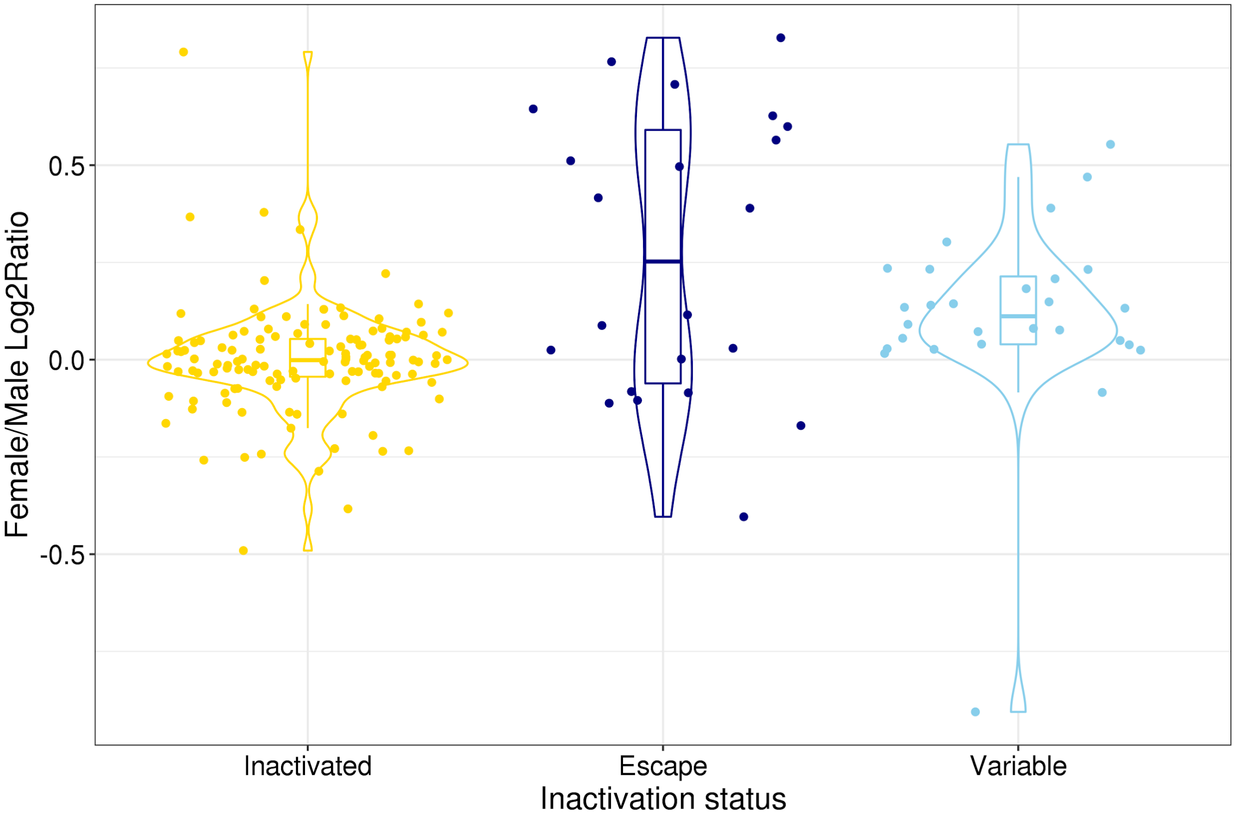

Figure S12. Total count and biased allele count for variants on genes that show opposite XCI patterns between the placenta and adult GTEx tissues and between the placenta.

For each gene, for each heterozygous and expressed variants, purple bars represent total RNA read count and light blue bars represent RNA read count of the biased allele. We observed that the total RNA read count for these variants are all greater than 10, suggesting that the patterns observed in Figure 4 is not due to technical artifacts.

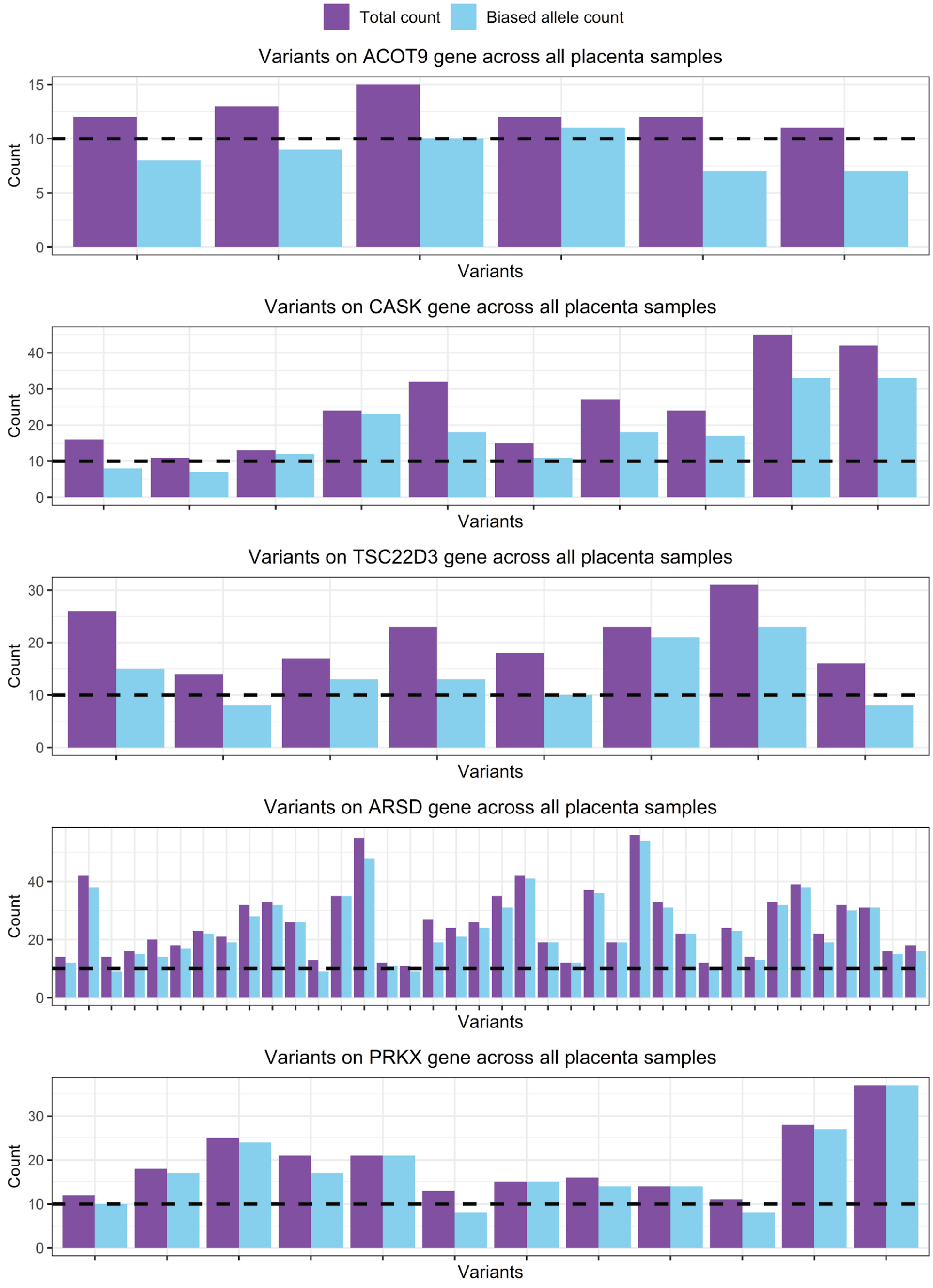

### Supplementary Notes

#### Note 1. Method to classify genes into genes that are inactivated, genes that escape XCI, and genes that show variable escape

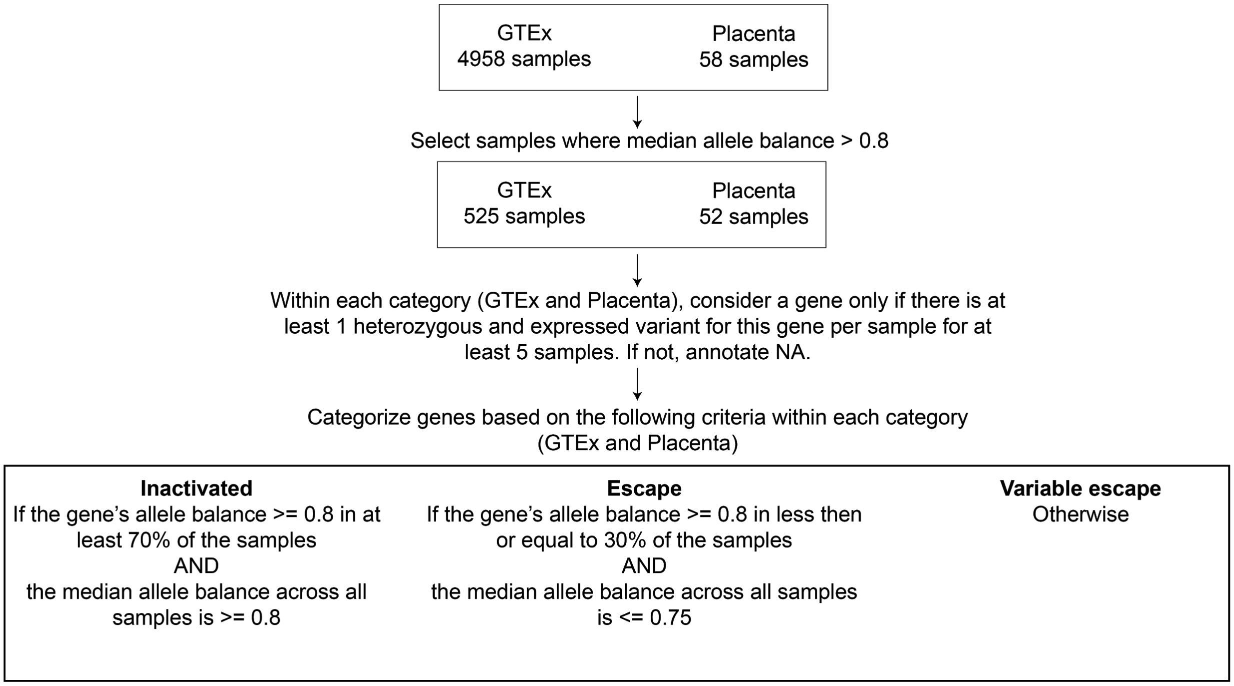
